# Individual differences in time-varying and stationary brain connectivity during movie watching from childhood to early adulthood: age, sex, and behavioral associations

**DOI:** 10.1101/2023.01.30.526311

**Authors:** Xin Di, Ting Xu, Lucina Q. Uddin, Bharat B. Biswal

## Abstract

Spatially remote brain regions exhibit dynamic functional interactions across various task conditions. While time-varying functional connectivity during movie watching shows sensitivity to movie content, stationary functional connectivity remains relatively stable across videos. These findings suggest that dynamic and stationary functional interactions may represent different aspects of brain function. However, the relationship between individual differences in time-varying and stationary connectivity and behavioral phenotypes remains elusive. To address this gap, we analyzed an open-access functional MRI dataset comprising participants aged 5 to 22 years, who watched two cartoon movie clips. We calculated regional brain activity, time-varying connectivity, and stationary connectivity, examining associations with age, sex, and behavioral assessments. Model comparison revealed that time-varying connectivity was more sensitive to age and sex effects compared with stationary connectivity. The preferred age models exhibited quadratic log age or quadratic age effects, indicative of inverted-U shaped developmental patterns. In addition, females showed higher consistency in regional brain activity and time-varying connectivity than males. However, in terms of behavioral predictions, only stationary connectivity demonstrated the ability to predict full-scale intelligence quotient. These findings suggest that individual differences in time-varying and stationary connectivity may capture distinct aspects of behavioral phenotypes.

## 1. Introduction

The interplay between distant brain regions is considered to be critical for comprehending brain functions. Functional connectivity is characterized by the statistical dependency between observed brain signals (Friston, 1994). This stationary characterization of functional connectivity is often explored during resting-state conditions (Biswal et al., 1995, 2010), providing valuable insights into the organizations of brain functional (Biswal et al., 2010; Margulies et al., 2016; Yeo et al., 2011). However, it is important to recognize that functional connectivity is not static but exhibit notable dynamics (Allen et al., 2014). The dynamic connectivity patterns across the entire brain encompass distinct “states” (Allen et al., 2014), demonstrating reliability (Abrol et al., 2017). Disruptions in dynamic connectivity have been linked to varies mental disorders (Fu et al., 2019).

Recently, movie watching has emerged as an alternative paradigm that bridges unconstrained resting-state conditions and well controlled task experiments. Watching movies allows participants to engage in a more “natural” experience than performing cognitive tasks. In addition, movie watching offers advantages over resting-state approaches in terms of enhanced compliance during scanning and potentially reduced artifacts resulting from head motion (Vanderwal et al., 2019). Interestingly, when individuals watch the same movie clip, their brain activity tends to exhibit similar patterns (Hasson et al., 2004), suggesting that these patterns hold functional significance. Additionally, time-varying connectivity measures demonstrates remarkable constancy across participants (Di et al., 2022b; Di and Biswal, 2020), further supporting the functional relevance of such time-varying measures of functional connectivity.

Time-varying and stationary functional connectivity are believed to capture different aspects of brain function. Many studies have revealed that the stationary connectivity patterns observed during the viewing of different movies exhibit remarkable similarity (Di et al., 2022b; O’Connor et al., 2017; Tian et al., 2021), and may even display high correlation with other mental states, such as resting-state (O’Connor et al., 2017). On the contrary, time-varying connectivity appears to depend on specific content of the movies, leading to substantial variations in dynamic patterns and the regions involved across different movie clips (Di et al., 2022b). This suggests that the time-varying connectivity may be more sensitive to reflect moment-to-moment brain function. More broadly, during the resting-state, time-varying connectivity has the ability to capture unique behavioral variability compared with stationary connectivity (Eichenbaum et al., 2021). Several studies focusing on disease classifications have shown that time-varying connectivity during the resting-state exhibits superior predictive power over stationary connectivity in distinguishing conditions such as schizophrenia, bipolar disorder (Rashid et al., 2016), and post-traumatic stress disorder (Jin et al., 2017).

To delve deeper into the functional implications of time-varying and stationary connectivity, our objective is to examine the individual differences in these connectivity measures during movie watching. Age and biological sex are common factors that contribute to variations among individuals. When watching movie clips, adults tend to exhibit higher levels of synchronized regional activity compared with children (Cantlon and Li, 2013; Petroni et al., 2018). However, children may display distinct response patterns compared with adults (Di and Biswal, 2022). A few studies have examined the impact of age on time-varying and stationary connectivity in the resting-state (Faghiri et al., 2018; Marusak et al., 2017; Rashid et al., 2018), revealing linear correlations between certain dynamic connectivity measures and age. In the current study, we employed more sophisticated age models and a model comparison framework to examine the age effects. We sought to determine whether time-varying and stationary connectivity differently portry age and sex effects.

Moreover, we inquire whether individual differences in time-varying and stationary connectivity are linked to behavioral outcome measures. Previous studies have demonstrated that, during the resting-state, time-varying connectivity exhibits superior predictive performance than stationary connectivity in relation to behavioral phenotypes (Eichenbaum et al., 2021) and the classification of mental disorders (Jin et al., 2017; Rashid et al., 2016). Therefore, it is reasonable to anticipate that time-varying connectivity during movie watching might also outperform stationary connectivity in predicting behavioral outcome measures.

In the current study, we conducted an analysis using fMRI data obtained from the Healthy Brain Network (HBN) (Alexander et al., 2017), which is a large open-access dataset. Previous studies have utilized this dataset to explore model-based brain activations (Richardson, 2019), stationary connectivity (Vanderwal et al., 2021), and event segmentation (Cohen et al., 2022) during the movie watching. The project involved the recruitment of female and male participants aged between 5 to 22 years, who underwent comprehensive behavioral assessments and MRI scanning. From the fMRI data during movie watching, we computed regional activity, stationary connectivity, and time-varying connectivity. To investigate the effects of age and sex on these brain measures, we employed a model comparison framework. Additionally, we employed a predictive modeling approach to examine the prediction capabilities of these brain measures in relation to behavioral outcomes. Our hypothesis posits that time-varying connectivity will exhibit stronger evidence of age and sex effects compared with stationary connectivity and regional activity. Furthermore, we anticipate that time-varying connectivity will outperform stationary connectivity and regional activity in terms of its predictive power for behavioral outcomes.

## 2. Materials and Methods

### 2.1. Healthy Brain Network dataset

#### 2.1.1. Dataset and participants

The MRI data utilized in this study were obtained from the Healthy Brain Network project website (http://fcon_1000.projects.nitrc.org/indi/cmi_healthy_brain_network/) (Alexander et al., 2017). We identified a total of 279 participants who have no diagnosis of any psychiatric or neurological disorders and have T1-weighted MRI data available (up to Release 9). Rigorous quality control measures were applied to both the T1-weighted structural images and fMRI images (details provided below). Among the participants, 159 individuals had structural images with motion artifacts or lesions. Additionally, participants with excessive head motion during fMRI scans (maximal framewise displacement smaller than one voxel) were further excluded. This led to the inclusion of 87 participants for ‘The Present’ dataset and 83 participants for the ‘Despicable Me’ dataset in the current analysis (Supplementary Figure S1). In total, 104 participants were included, with 66 individuals having watched both video clips. Among the included participants, there were 61 males and 43 females, with ages ranging 5.0 to 21.9 years (*Mean* = *12.0*; *Standard Deviation* = *4.1*). The participants’ family socioeconomic status was approximated using the Barratt Simplified Measure of Social Status (BSMSS) (http://socialclassoncampus.blogspot.com/2012/06/barratt-simplified-measure-of-social.html).

#### 2.1.2. MRI data

We conducted an analysis of fMRI data collected while the participants were engaged in watching two animated movie clips. The first clip was a short film titled ‘The Present’ (duration: 3 minutes and 21 seconds, Filmakademie Baden-Wuerttemberg, 2014), while the second clip consisted of a 10-minute segment extracted from the animated film ‘Despicable Me’ (Illumination, 2010). Additionally, the high-resolution anatomical MRI images were used for preprocessing purposes.

The MRI data were acquired from two MRI centers, Rutgers University Brain Imaging Center (RUBIC) with a 3T Siemens Trio scanner, and Citigroup Biomedical Imaging Center (CBIC) with a 3T Siemens Prisma scanner. Out of the 104 participants included in the current analysis, 66 were scanned at the RUBIC site, and 38 were scanned from the CBIC site. The scanning protocols were similar between the two sites. For the fMRI scans, the key imaging parameters were as follows: TR = 800 ms; TE = 30 ms; flip angle, 31°; voxel size = 2.4 x 2.4 x 2.4 mm^3^; multi-band acceleration factor = 6. As for the T1-weighted anatomical MRI scans, the images were acquired using either the Human Connectome Project (HCP) or the Adolescent Brain Cognitive Development (ABCD) sequences. The sequences differ in terms of voxel sizes; however, the anatomical images were solely used for preprocessing of the fMRI images. For more detailed information about the MRI protocols, please refer to the HBN project website and (Alexander et al., 2017).

#### 2.1.3. Behavioral measures

We picked two behavioral measures, full-scale intelligence quotient (FSIQ) from the Wechsler Intelligence Scale for Children – Fifth Edition (Wechsler, 2014) and the Social Communication Questionnaire (SCQ) (Rutter et al., 2003). FSIQ serves as an indicator of general cognitive ability and is commonly employed in studies investigating the relationship between brain function and behavior (Vieira et al., 2022). Among the participants who watched the video clip ‘The Present’, FSIQ scores were available for 64 individuals, while for the clip ‘Despicable Me’, FSIQ scores were available for 60 participants. On the other hand, the SCQ is a parent-report questionnaire that measures social and communication symptoms related to autism spectrum disorder. In a previous study utilizing the same dataset, it was reported that the SCQ scores were associated with brain activation during specific time points (events) (Richardson, 2019). For the video clip ‘The Present’. SCQ scores were available for 70 participants, and for the clip ‘Despicable Me’, SCQ scores were available for 65 participants.

### 2.2. MRI data processing

#### 2.2.1. Structural MRI quality control and processing

Following the established protocol employed in our laboratory (Di and Biswal, 2023), we performed pre-processing and quality control procedures on the MRI data. A thorough visual inspection was conducted on the MRI images of all participants to identify any issues such as excessive head motion, partial coverage, or brain lesions. Both the T1-weighted and segmented images underwent visual quality control assessment. Out of the 279 total participants, 159 individuals had images exhibiting ghost artifacts, motion artifacts, or lesions. Consequently, 120 participants were selected for further analysis. Statistical parametric mapping (SPM12, https://www.fil.ion.ucl.ac.uk/spm/) in MATLAB (R2021a, https://www.mathworks.com/) was used for MRI image processing. The T1-weighted image for each participant underwent segmentation into various tissue types, including gray matter, white matter, cerebrospinal fluid, and others. The segmented images were then roughly aligned to the standard Montreal Neurological Institute (MNI) space using linear transformation. Then the DARTEL procedure was used to register the segmented gray matter and white images across all the individuals and generate a sample-specific template through multiple iterations (Ashburner, 2007). The resulting averaged gray matter template was then linearly normalized to MNI space.

#### 2.2.2. Functional images preprocessing

The functional images were realigned to the first image, coregistered to the anatomical image, and then normalized into the MNI space. During the normalization step, the functional images were resampled into a voxel size of 2.4 x 2.4 x 2.4 mm^3^, and spatially smoothed using an 8 mm Gaussian kernel. Lastly, a voxel-wise general linear model (GLM) was used to remove head motion artifacts and low-frequency drifts. The GLM included Friston’s 24 head motion parameters (Friston et al., 1996) and a high-pass filter at 1/128 Hz. The residual images obtained from the GLM step were used for further analysis. To ensure accuracy of each preprocessing step, a visual inspection was carried out following each step (Di and Biswal, 2023).

Head motion is an important factor that can affect BOLD fMRI signals. To address this, we excluded participants whose maximum framewise displacement exceeded 2.4 mm or 2.4° (approximately the size of a voxel) in any direction or during any movie clip. Next, we examined the age effects on head motion. First, we showed that the framewise displacement time series were not synchronized across participants, as indicated by the first principal component (PC) explaining less than 5% of the variance (Supplementary Figure S2A). Secondly, we used the mean framewise displacements in translation and rotation as measures of head motion and examined their age effects. Model comparison analyses showed that, for the ‘The Present’ clip, a constant model without age effects was favorable (Supplementary Figure S2C and S2D). However, for the ‘Despicable Me’ clip, there was evidence of log age effects (Supplementary Figure S2E and S2F). Notably, the age-related patterns observed in head motion differ significantly from those observed in the brain measures. Nevertheless, we added the mean framewise displacement of translation and rotation into the age fitting models.

#### 2.2.4. Independent component analysis

We employed independent component analysis (ICA) to reduce the dimensionality of the fMRI data (Di et al., 2022b; Di and Biswal, 2022). The ICA was performed using the Group ICA Of fMRI Toolbox(GIFT) (Calhoun et al., 2001) with data from both video clips combined. A total of twenty independent components (ICs) were extracted and visually inspected. Among them, 18 components were considered functional meaningful networks based on our previous work (Di et al., 2022b; Di and Biswal, 2022). Four networks are of particular interest due their involvement in movie watching (Di et al., 2022b; Di and Biswal, 2022): the dorsal visual network, temporoparietal junction, supramarginal network, and the default mode network (particularly the posterior cingulate cortex) (Figure 2A). For each participant and video clip, the time series of the 18 networks were back reconstructed and utilized for further analysis.

**Figure 1.**
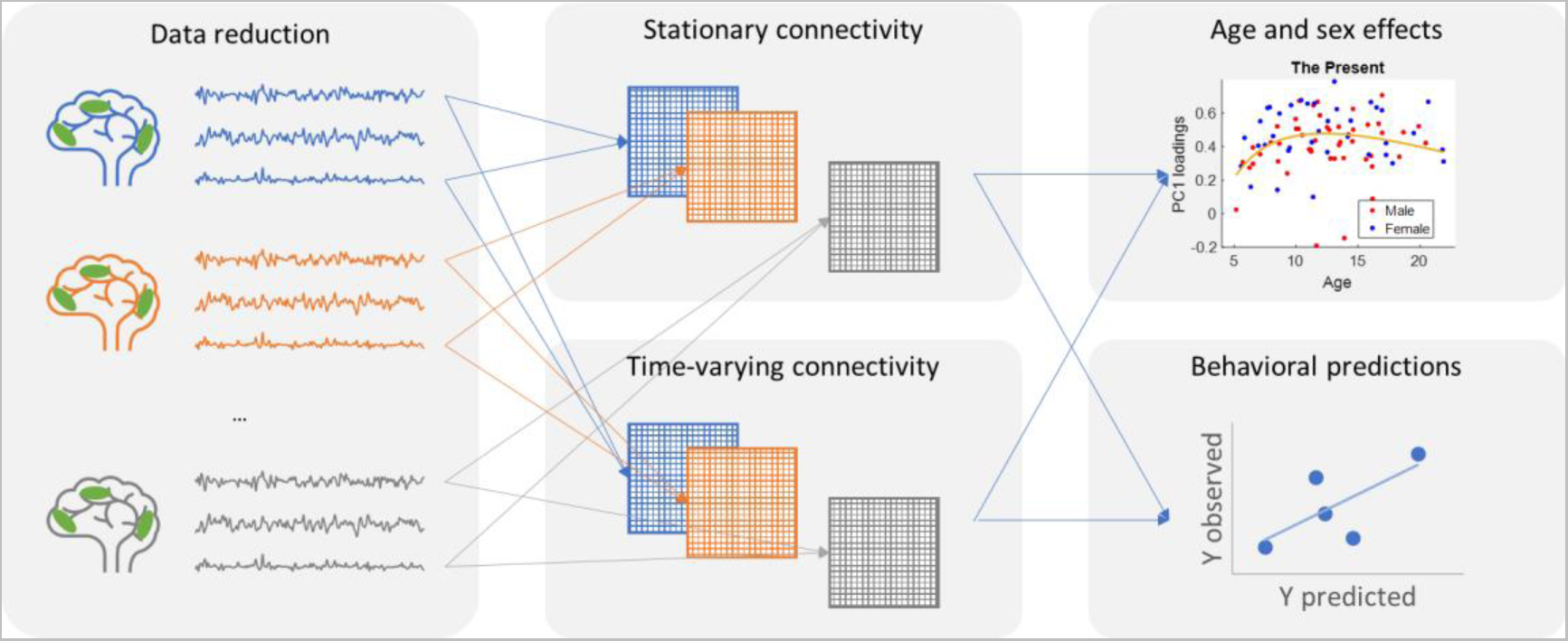
Summary of main data analysis steps in the current study. After preprocessing independent component analysis (ICA) was applied to the fMRI data to reduce the data dimension to 19 spatial networks and the associated time series for each participant and movie clip. Stationary and time-varying connectivity matrices were calculated for each individual and movie clip. Age and sex effects on connectivity measures were then studied using a model comparison approach. Lastly, machine learning regression was used to study behavioral associations with the different connectivity measures.

**Figure 2.**
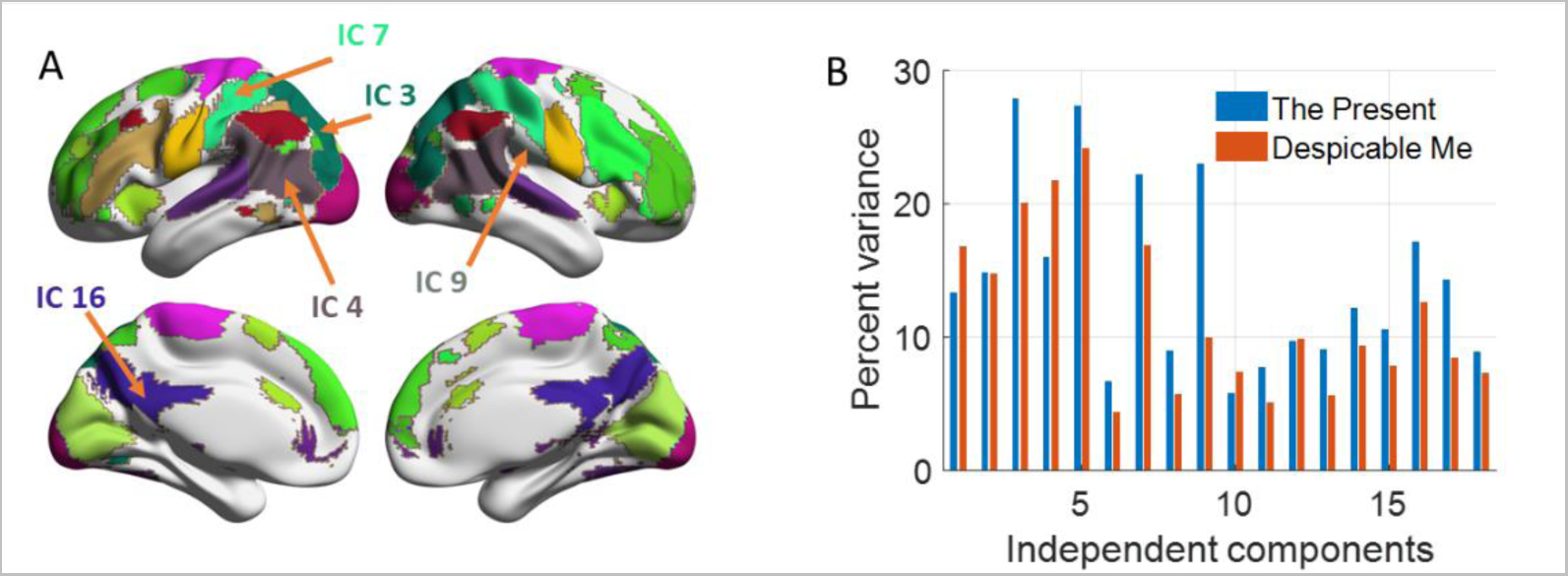
A, Maps of eighteen independent components that were included in the current analysis. The maps were thresholded at z > 3 after z transformations of the original IC maps, and were shown in a winner-take-all manner when overlapping. The arrows indicate the networks of interest for movie watching, IC3, dorsal visual; IC4, temporoparietal junction; IC7 supramarginal; IC9, secondary somatosensory; and IC16, posterior cingulate. BrainNet Viewer was used for visualization (Xia et al., 2013). B, inter-individual consistency of regional activity (percent variance explained by the first principal component) for the two video clips.

ICA is a data-driven method that extracts sources of brain activity based on the current sample and experimental design, which might limit the generalizability of the current results. To illustrate the robustness of the current results, we adopted an independent ICA template and replicated all the statistical analyses. The template maps of 16 ICs were derived from a separate dataset of movie-watching experiments in our previous study (Di et al., 2022b). These IC maps were resampled into the voxel dimension of the current preprocessed fMRI data. Subsequently, the IC maps were regressed against the current preprocessed fMRI data, resulting in 16 time series for each participant and movie clip. This approach is akin to the first stage of a dual-regression analysis (Nickerson et al., 2017).

### 2.3. Statistical analysis

#### 2.3.1. Regional activity and connectivity measures

Inter-subject correlation has been used to index shared responses during movie watching (Hasson et al., 2004; Nastase et al., 2019). In this study, we used a principal component analysis (PCA) based method to estimate inter-individual consistency (Di and Biswal, 2022). For each network (IC), the time series data from each participant formed a matrix of dimensions *t* x *n*, where *t* and *n* represent the number of time points and participants, respectively. The matrices were of size *250* x *87* for the clip ‘The Present’, and *750* x *83* for the clip ‘Despicable Me’. We performed PCA on the matrix, and obtained the variances explained by the first and second PCs. To establish a null distribution, a circular time-shift randomization method was used with 10,000 randomizations (Di and Biswal, 2022; Kauppi et al., 2010). Because none of the second PCs explained statistically significant variance, only the first PCs were used for the analysis of individual differences.

To assess time-varying connectivity between each pair of two networks (ICs), we calculated point-by-point interactions (multiplications) as an index (Di et al., 2022b; Faskowitz et al., 2020). Subsequently, PCA was employed to estimate the inter-individual consistency in the time-varying connectivity. The loadings of the first PC were used as a measure of individual differences in time-varying connectivity.

Lastly, we calculated stationary connectivity as the Pearson’s correlation between the time series of each pair of the 18 networks (ICs).

#### 2.3.2. Age and sex effects

We adopted a model comparison framework to examine age and sex effects on regional activity, stationary connectivity, and time-varying connectivity. Specifically, we constructed five models to assess the age effects on each brain measure, while considering sex as a separate regressor. Additional covariates included a scanner site variable and mean framewise displacement in translation and rotation. The five models are as follows,

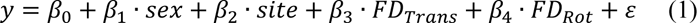

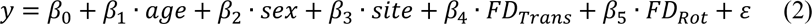

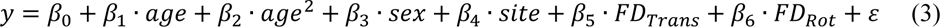

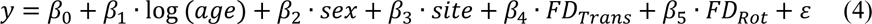

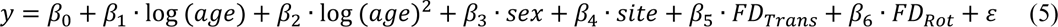

Where y represents a specific measure such as regional activity in a network, stationary connectivity, or time-varying connectivity between two networks. Model 1 represents a baseline condition where there is no age effect. Models 2 and 3 represent linear age effect and quadratic age effect models. Models 4 and 5 represent log age effect and quadratic log age effect models. The log age models consider the fact that brain measures may grow faster and then decrease slower within the studied age range. We additionally built five models the same as models 1 through 5 except that there were no sex effects in each of the models. Therefore, we had 10 models in total (2 x 5).

To compare different age models and sex effects, we used a model comparison procedure. A total of 10 models were fitted using ordinary least square, and the Akaike information criterion (AIC) was calculated. We then calculated Akaike weights (Wagenmakers and Farrell, 2004) for each model, which quantify the model evidence relative to the best model among all the 10 models, with the sum of all the models as 1. The Akaike weights of the same age model with and without the sex term were added as the model evidence of a particular age effect, regardless of the sex effects (Portet, 2020). This allowed us to examine the relative strength of different age models in explaining the observed developmental effects. Similarly, for the sex effect, we added the Akaike weights for all five age models with the sex term. The cumulative model weights depend on the number of alternative models. We adopt a threshold of 0.6 for the age model comparison (involving 5 models) and 0.8 for the sex effect comparison (involving 2 models).

#### 2.3.3. Behavioral prediction analysis

We applied ridge regression to study brain-behavioral associations. The predicted variable, either FSIQ scores or SCQ scores, was represented as an *n* by *1* vector. *N* varied depending on the availability of data for FSIQ, SCQ, and the two movie clips. The predicting variables were either regional activity, stationary connectivity, or time-varying connectivity. We applied a leave-one-out cross-validation to evaluate the prediction performance of each brain measure. Specifically, we held out one participant’s data, and used the remaining *n – 1* data to obtain a prediction model. We used a linear model for the prediction.

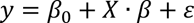

where *y* is a *n – 1* vector of either FSIQ of SCQ scores, *X* is a *n – 1* by *m* matrix of regional activity, stationary connectivity, or time-varying connectivity matrices. For the regional activity, *m* equals to 18 of the networks. For the connectivity matrices, the number of column equals *153* (*18 x 17 / 2*), which was larger than the number of rows. To align the dimensions, a dimension reduction procedure was applied to select the top 18 features with the highest absolute correlations with the predicted variable. Therefore, *X* always had dimensions of *n – 1* by *18*.

The model was fitted with a ridge regularization. The regularization parameter *λ* was determined using a nested cross-validation procedure for each training set. With the optimal *λ*, the model was trained using the *n – 1* training data. The model was applied to the held-out participant to calculate the predicted value. The procedure was performed *n* times for the *n* participants, resulting in *n* predicted values. We calculated the correlation between the predicted value and the actual values across all n participants to obtain an estimate of prediction accuracy.

The optimal *λ* was determined for each leave-one-out sample using an inner leave-one-out loop. Within the *n – 1* outer-loop training set, we built linear models with *n – 2* individuals with 21 *λ* values (from *2^-10^* to *2^0^* in logarithmical space). The prediction accuracies across all the inner loop samples were calculated for all the *λ* values. The *λ* with the highest correlation was parsed to the outer loop as the optimal *λ* for model training and prediction. The prediction procedure is outlined in Supplementary Figure S3.

The prediction accuracies were assessed using the correlation coefficient between the predicted and observed values. In total, there were 12 predictions, including 2 movies, 3 brain measures, and 2 behavioral measures. We used false discovery rate correction to account for multiple comparisons (q < 0.05).

## 3. Results

### 3.1. Characteristics of the included sample

We implemented a rigorous quality control protocol for both structural and functional MRI images, resulting in exclusion of a substantial number of participants. To assess potential differences between the included and excluded samples, we compared their demographic characteristics, as summarized in Table 1. The analysis revealed no significant differences in the distributions of biological sex (*χ^2^ = 0.0162, p = 0.898*). However, the excluded group exhibited a significantly younger age than the included group (*t = 6.54, p < 0.001*). Furthermore, within the participants with available data, we examined the differences in intelligence (FSIQ) and social economic status (BSMSS) between the two groups. No statistical differences were observed for either FSIQ (*<u>t = 1.68, p = 0.094</u>*<u>) or</u> BSMSS (*t = 0.83, p = 0.41*).

**Table 1.**
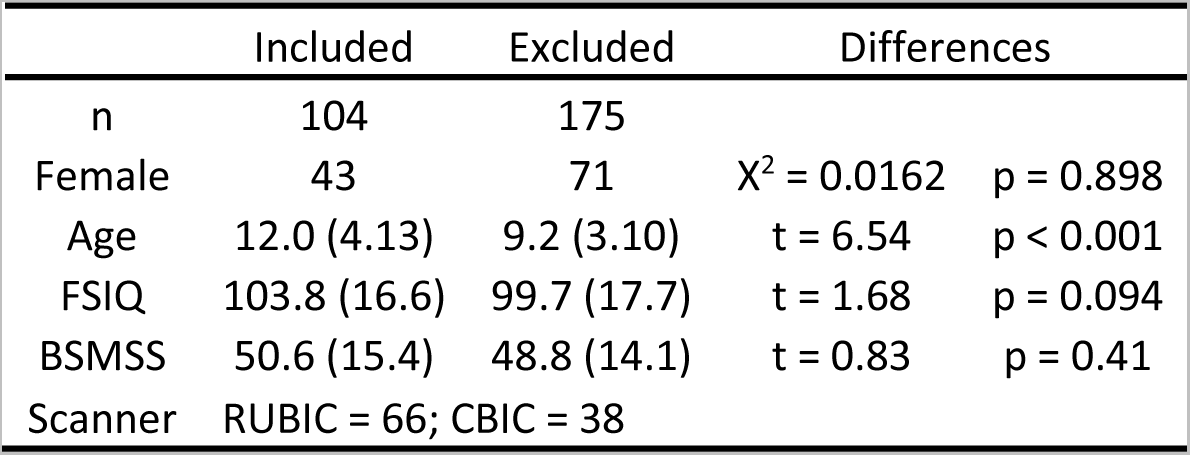
Demographic information for the participants included and excluded from the current analysis. Notably, not all participants completed the FSIQ and BSMSS assessments. Specifically, among the included and excluded groups, FSIQ score was available for 78 and 146 participants, respectively, while BSMSS scores were available for 80 and 129 participants, respectively.

|  | Included | Excluded | Differences |  |
| --- | --- | --- | --- | --- |
| n | 104 | 175 |  |  |
| Female | 43 | 71 | $\chi^2 = 0.0162$ | $p = 0.898$ |
| Age | 12.0 (4.13) | 9.2 (3.10) | $t = 6.54$ | $p < 0.001$ |
| FSIQ | 103.8 (16.6) | 99.7 (17.7) | $t = 1.68$ | $p = 0.094$ |
| BSMSS | 50.6 (15.4) | 48.8 (14.1) | $t = 0.83$ | $p = 0.41$ |
| Scanner | RUBIC = 66; CBIC = 38 |  |  |  |

### 3.2. Regional activity

We performed PCA to assess the extent to which shared brain activity could be explained by a single or multiple components. For each of the 18 networks (ICs), the first PC exhibited a statistically significant amount of variance explained for both videos, as determined by circular time-shift randomization testing. Conversely, the second PCs in all the networks didn’t account for a significant amount of variance. Therefore, we focused solely on the first PCs in the following analysis. Figure 2B shows the percentage variance explained by the first PC in the 18 networks for the two movie clips. In addition to lower-level sensory networks such as the visual and auditory networks, several higher-level networks showed substantial inter-individual consistency, including the dorsal visual network (IC3), temporoparietal junction network (IC4), supramarginal network (IC7), and default mode network (IC16). There are also noticeable differences between the two video clips. In particular, a network covering the posterior insula, secondary somatosensory regions and cingulate (IC9) showed more than twice the variance explained by the first PC in ‘The Present’ (23.0%) than ‘Despicable Me’ (10.0%). To gain insights into the cognitive implications of this network, we submitted the map to cognitive decoding using Neurosynth (Yarkoni et al., 2011). After removing terms related to brain labels, the top five terms associated with this network were pain, painful, tactile, stimulation, and touch.

We then applied a model comparison procedure to examine the age and sex effects. Figure 3A and 3B depicts the model evidence for the five age models across the 18 network ICs and the two video clips. Networks that exhibited a strong preference for a particular model typically favored the quadratic log age or quadratic age model. We identified the network ICs with model evidence exceeding 0.6 for a specific age model and plotted the fitted effects in Figure 3D and 3F. All the age effects showed an inverted-U shape, with peak around 10 years of age. The quadratic log age models indicated a steeper increase in the younger ages and slower decrease in older ages. Figure 3E and 3G further present individual loadings as well as the fitted curves for the two curves on the top of Figure 3D and 3F. Specifically, Figure 3E corresponds to the supramarginal network (IC7) in the video clips of ‘The Present’, and Figure 3G corresponds to the bilateral parietal junction network (IC4) in the ‘Despicable Me’ clip.

**Figure 3.**
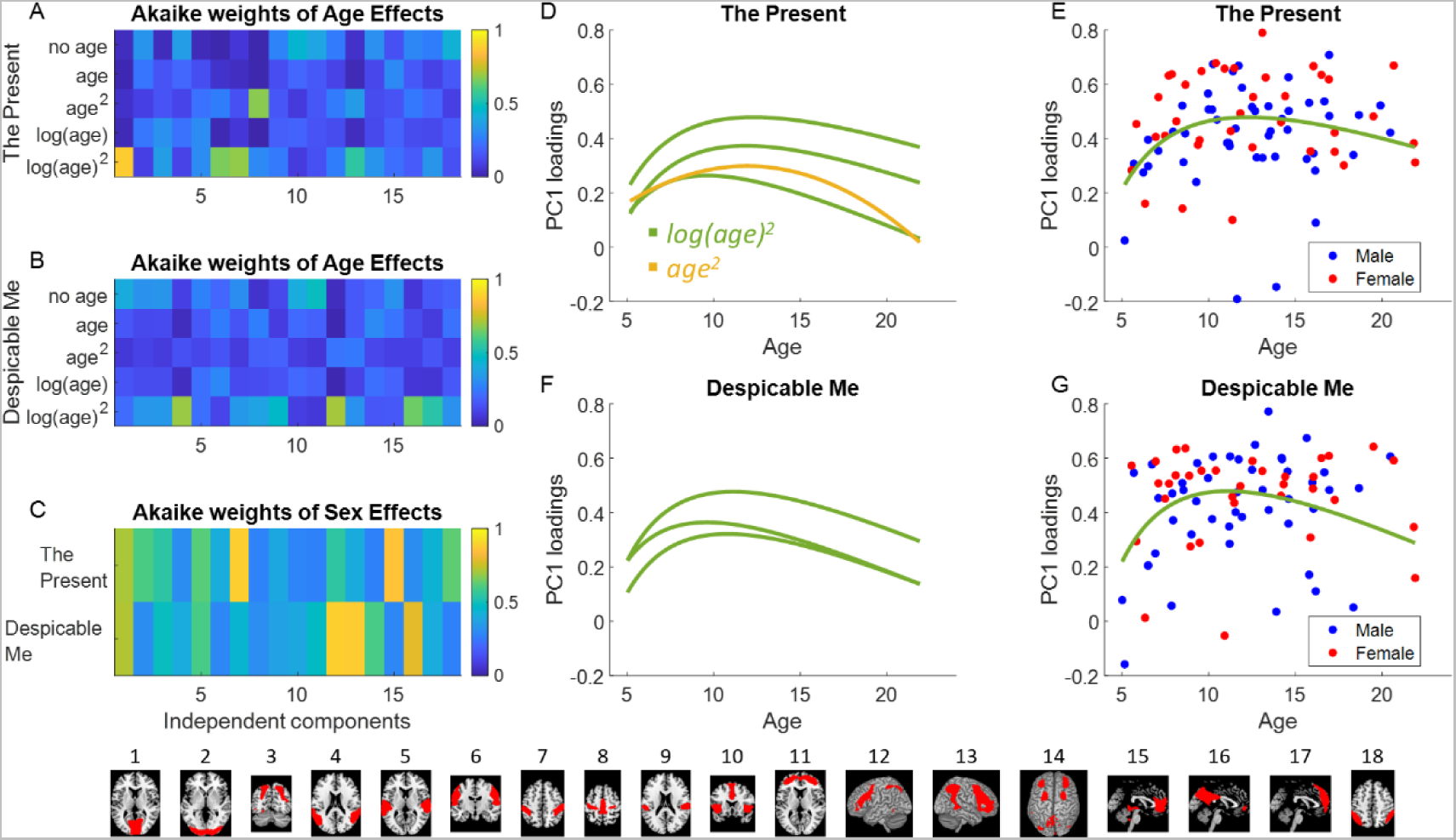
Akaike weights for different age models for regional activity in the 18 networks (independent components) for the video clips “The Present” (A) and “Despicable Me” (B). C shows the model evidence for the sex effects. D and F show fitted age effects in networks that exhibited Akaike weights greater than 0.6 in a specific model. E and G show the fitted age curves and individual scatter plot in two representative networks, corresponding to the top curves in D and F, respectively.Figure 4 A and E, group averaged stationary connectivity among 18 independent component (IC) networks for the two video clips. B and F, connectivity with winning age models with model evidence greater than 0.6. C and G, fitted curves with corresponding color representing the specific age models. D and H, connectivity with evidence of sex effects greater than 0.8. The bottom row shows the representative maps of the 18 networks.

Four networks showed strong evidence of a sex effect (> 0.8) on regional activity, which varied between the two video clips (Figure 3C). For ‘The Present’, the supramarginal network (IC7) (model probability = *88.36%*) and medial frontal network (IC15) (model probability = *85.76%*) showed evidence of sex effects. While for the video clip of ‘Despicable Me’, the left (IC12) and right (IC13) fronto-parietal networks showed evidence of sex effects (model probability = *88.80%* and *87.50%*, respectively). In all cases, females exhibited higher consistency than males.

### 3.3. Stationary connectivity

The group averaged stationary connectivity matrices for the two video clips are shown in Figure 4A and 4E, respectively. The two matrices exhibited a similarity, consistent with our previous findings (Di et al., 2022b). The higher levels of stationary connectivity primarily occur between networks with similar functions, e.g., among visual networks and among fronto-parietal networks.

**Figure 4.**
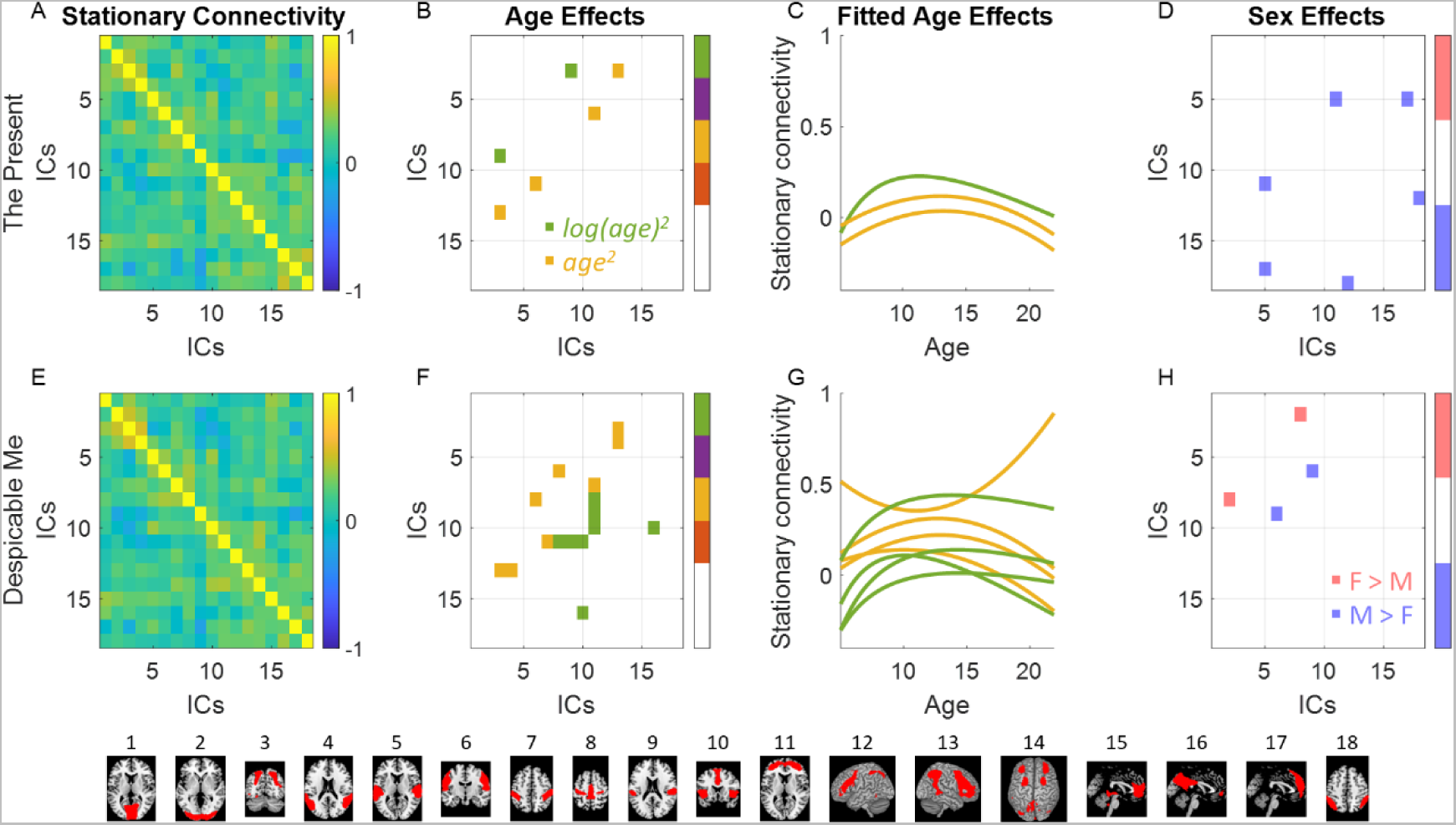
A and E, group averaged stationary connectivity among 18 independent component (IC) networks for the two video clips. B and F, connectivity with winning age models with model evidence greater than 0.6. C and G, fitted curves with corresponding color representing the specific age models. D and H, connectivity with evidence of sex effects greater than 0.8. The bottom row shows the representative maps of the 18 networks.

The full model comparison results for the age and sex effects on stationary are shown in Supplementary Figure S4. With a model evidence threshold of 0.6 in any specific age effects, the connections with significant age model preferences are shown in Figure 4B and 4F for the two video clips respectively. The preferred age models tended to be either quadratic log age or quadratic age models. An inverted-U shape was observed for most age effects, as shown in Figure 4C and 4G. Specifically, three connections showed a preference for an age model for the clip of ‘The Present’, while eight connections showed a preference of an age model for the ‘Despicable Me’ video. The significant age effects appeared to be distinct between the two video clips. Notably, one stationary connectivity during ‘The Present’ was particularly interesting, because it involved two networks of interest, i.e., the dorsal visual network (IC3) and secondary somatosensory network (IC9). Lastly, three connections for the clip of ‘The Present’ and two connections for the ‘Despicable Me’ clip showed sex effects, although they were not among the networks of interest specifically related to movie watching.

### 3.4. Time-varying connectivity

Figure 5A and 5E show the consistency of time-varying connectivity for the two movie clips. In general, the clips of ‘The Present’ showed higher consistency in time-varying connectivity. Interestingly, several many networks of interest, including the dorsal visual network (IC3), supramarginal network (IC7), and secondary somatosensory network (IC9) demonstrated high consistency in time-varying connectivity. Moreover, the medial prefrontal network (IC17), which is part of the default mode network, also showed consistent time-varying connectivity with the other regions of interest.

**Figure 5.**
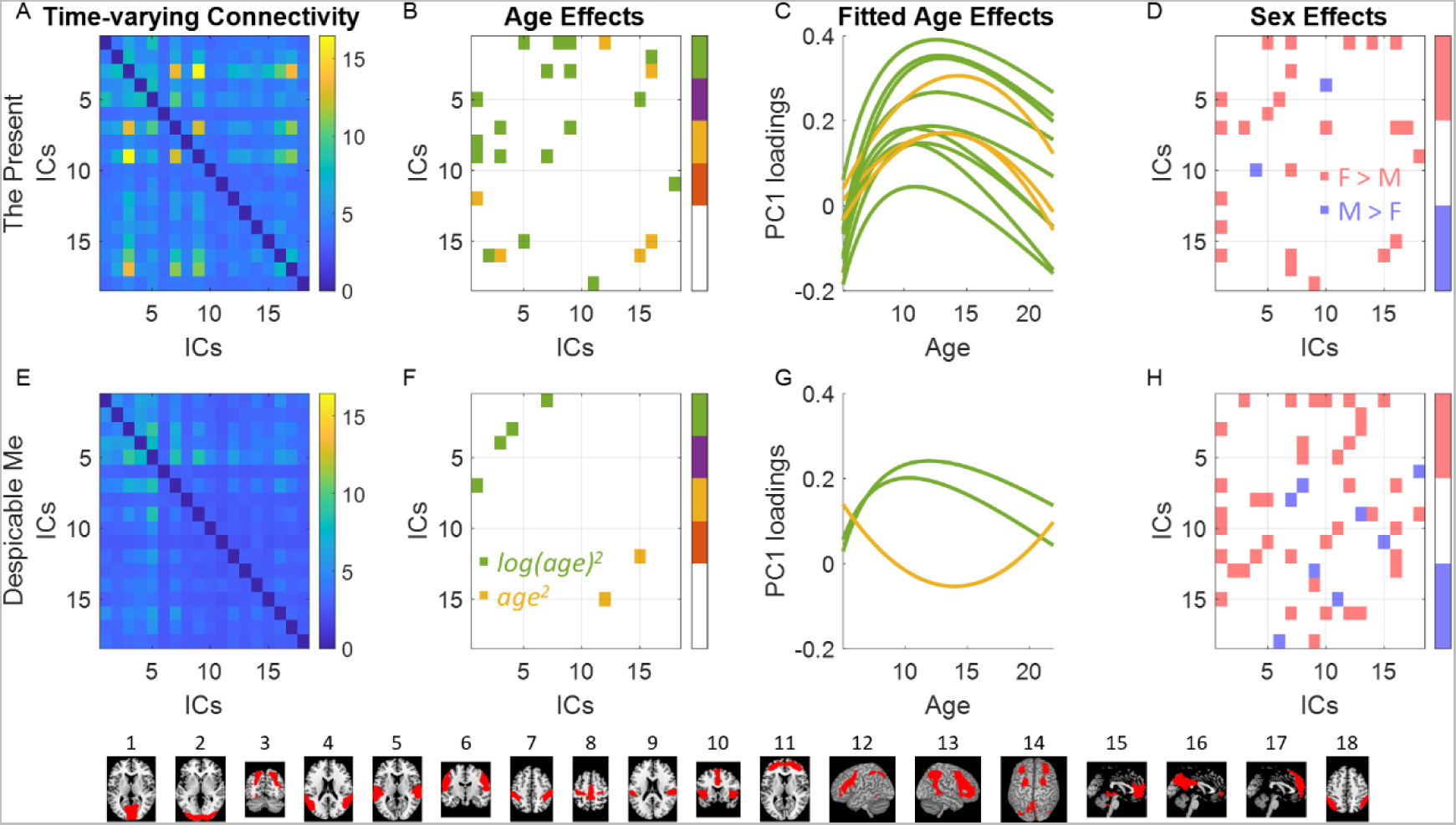
A and E, the inter-individual consistency of time-varying connectivity (percent variance explained by the first principal component) among 18 networks (independent components, ICs) for the two video clips. B and F, connectivity with winning age models with model evidence greater than 0.6. C and G, fitted curves with corresponding color representing the specific age models. D and H, connectivity with evidence of sex effects greater than 0.8. The bottom row shows the representative maps of the 18 networks.

The age effects on time-varying connectivity also tended to favor quadratic log age or quadratic age effects, as shown in Supplementary Figure S5, and Figure 5B, and 5F. Fourteen connections showed strong preferences for an age model in the video ‘The Present’. Interestingly, time-varying connectivity among the dorsal visual (IC3), supramarginal (IC7), and secondary somatosensory (IC9) networks strongly preferred the quadratic log age effects. In contrast, only three connections showed strong preferences to an age model in the video ‘Despicable Me’.

Many connections also showed strong evidence of sex effects, mainly indicating higher consistency in females than males (Figure 5D and 5H). This included the time-varying connectivity between the dorsal visual (IC3) and the supramarginal networks (IC7), which exhibited strong overall consistency in the video ‘The Present’.

### 3.5. Behavioral relevance

We employed machine learning regression to examine the behavioral relevance of regional activity, stationary connectivity, and time-varying connectivity. Our analyses focused on two behavioral measures: FSIQ and SCQ scores. Using a leave-one-out cross-validation, we estimated the prediction accuracies of the three brain measures for FSIQ and SCQ scores. Interestingly, only stationary connectivity showed statistically significant prediction accuracies (Figure 6). By using stationary connectivity from both video clips, we achieved prediction accuracies of around 0.35, as indicated by the correlations between predicted and actual FSIQ scores. However, individual differences in regional activity or time-varying connectivity did not yield predictive power for FSIQ scores (Supplementary Figure S6). In addition, none of the brain measures were able to predict SCQ scores (Supplementary Figure S7).

**Figure 6.**
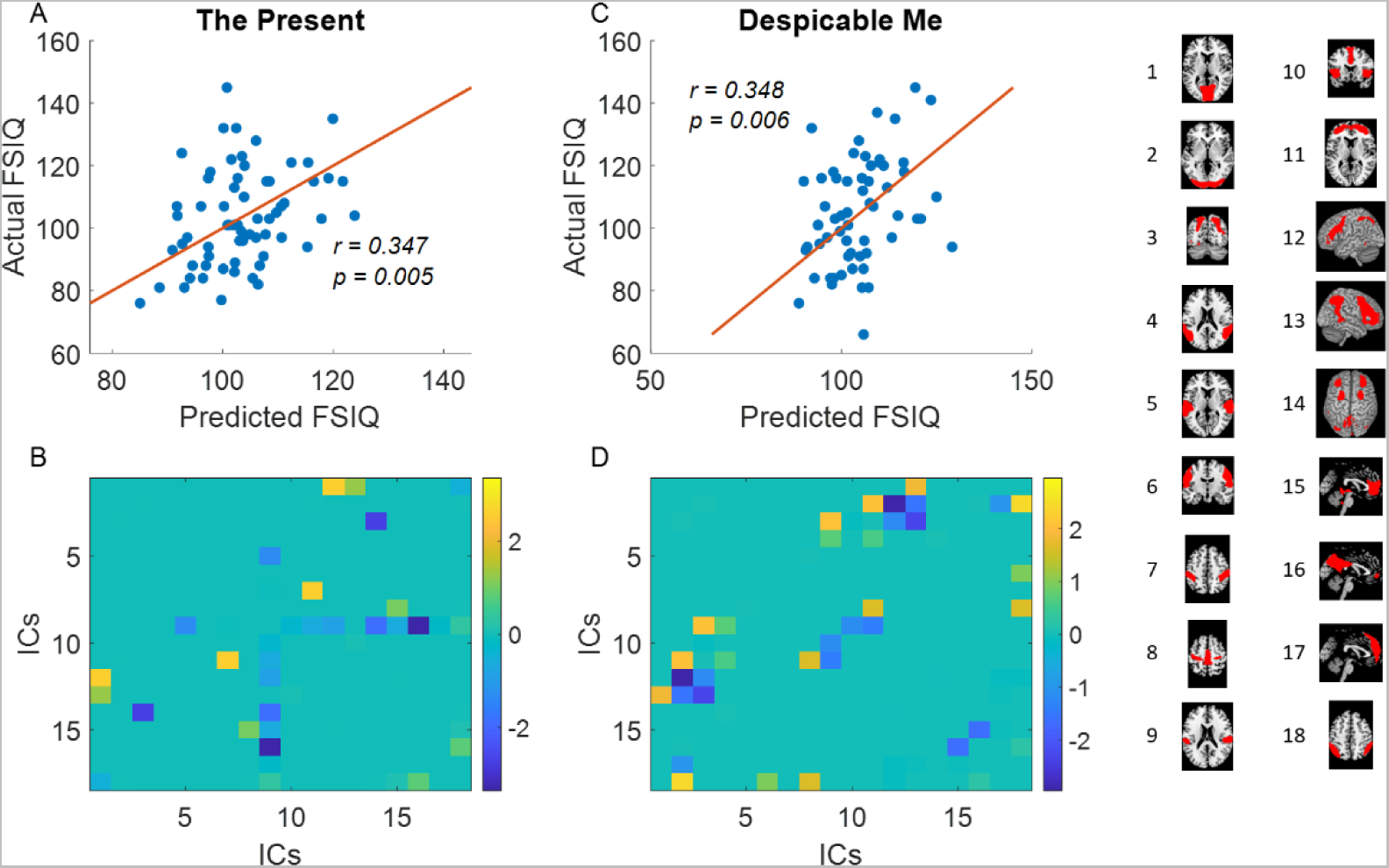
Results of full-scale intelligence quotient (FSIQ) predictions using stationary connectivity for the two video clips. Top row, each dot represents a predicted value using leave-one-out cross validation and its corresponding actual value. The red line indicates *y = x*. Bottom row, averaged weights of the prediction model across all the LOO models. The maps of the corresponding independent components (ICs) are shown on the right.

### 3.6. Replicability with independent IC maps

To assess the reliability of the analysis approach, we conducted a replication analysis using independent IC maps from our previous study (Di et al., 2022b). These 16 independent maps consisted of networks similar to those identified in the current dataset, including different visual networks, auditory network, and supramarginal networks (Figure 7). However, the secondary sensorimotor network (original IC9) was not present in the replication maps, indicating its specificity to the current dataset. The replication networks exhibited similar patterns of activation consistency as the original networks (Supplementary Figure S8). Regarding age effects, the preferred age models in the replication analysis were mostly log age square or age square models, which aligns with the main results. Furthermore, time-varying connectivity demonstrated greater sensitivity to age effects than stationary connectivity. However, the specific connections displaying age effects did not completely overlap with those observed in the main results. In terms of sex effects, the replication results provided more evidence of sex differences, despite including a smaller number of ICs than the main results. Females generally exhibited stronger stationary connectivity and higher levels of consistency in time-varying connectivity than males. Finally, concerning behavioral predictions, none of the brain measures or video clips yielded significant predictions for FSIQ and SCQ scores.

**Figure 7.**
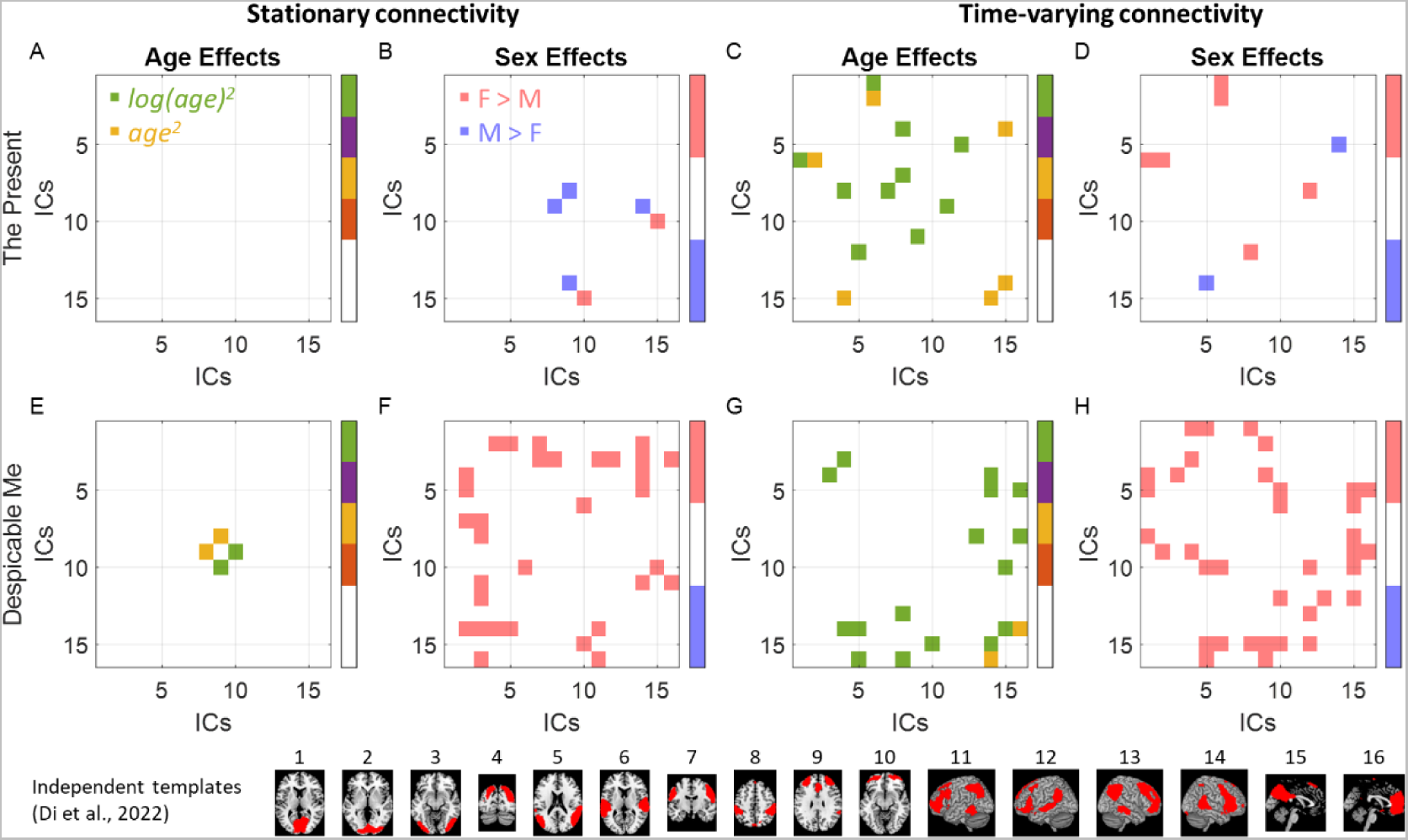
Replication analysis of age and sex effects on stationary connectivity and time-varying connectivity using independent templates from Di et al., (2022).

## 4. Discussion

In the current study, we examined individual differences in stationary connectivity and time-varying connectivity, as well as regional activity while participants watched movies, focusing on a sample of children to young adults. Consistent with our hypothesis, time-varying connectivity exhibited greater sensitivity to age and sex effects compared with stationary connectivity and regional activity. Contrary to our hypothesis, however, only stationary connectivity demonstrated predictive capabilities for FSIQ scores.

The two animated video clips evoked consistent brain activations across individuals in higher-order brain regions, including the dorsal visual, temporoparietal junction, supramarginal, and the default mode networks. These findings align with previous studies that have observed similar consistent responses with different movie clips and samples (Di et al., 2022b; Di and Biswal, 2022). Many of these regions, e.g., the temporoparietal junction, supramarginal, and default mode networks are known to be involve in processing of higher-order social information, which is expected during watching of the movie clips. More interestingly, a unique independent component encompassing the secondary somatosensory cortex, posterior insula, and cingulate cortex exhibited significantly higher consistency in the clip of ‘The Present’ compared with ‘Despicable Me’. These regions are associated with higher somatosensory and pain process, and may be involved in empathy for pain (Allen et al., 2017; Lamm et al., 2007). Given that ‘The Present’ includes a scene featuring an amputated limb, it is reasonable to observe the involvement of these regions. Our subsequent discussions will focus on these networks.

For all the connectivity and activity measures examined, the optimal age models consistently revealed quadratic log age or quadratic age effects, predominantly exhibiting an inverted-U shape. Time-varying and stationary connectivity showed age effects in different connections, with time-varying connectivity demonstrating age effects between regions related to movie watching. The presence of inverted-U shape indicates that the brain measures tend to increase during early childhood, reaching a peak, and subsequently decline towards adulthood. These findings provides a more comprehensive understanding of synchronized responses compared with prior studies that focused on only two groups of adults and children (Cantlon and Li, 2013; Petroni et al., 2018). The reduced synchrony in adults, compared with teen age children, may suggest that neural processing becomes more efficient with age, requiring less activation. Alternatively, it is possible that adult participants may exhibit more idiosyncratic responses to the video clips, or their engagement may be diminished due to the cartoon nature of the stimuli. Nevertheless, a practical implication of these findings is that when accounting for age effects, a simple linear model may not adequately capture the complexity of the relationship.

Several instances of connectivity exhibited a U-shaped pattern, suggesting elevated values in both younger and older age groups compared to adolescents. This observation was bolstered by reducing the model comparison threshold, revealing additional stationary connectivity displaying a similar U-shaped pattern. Previous studies have reported analogous patterns during resting-state, primarily within higher-order brain regions (Zuo et al., 2010). These findings likely reflect the dynamic developmental patterns of segregations and integrations throughout childhood (Cao et al., 2014; Zuo et al., 2010). But it is noteworthy that these connections exhibiting U-shaped age effects did not involve the brain networks of interest related to movie watching.

We observed widespread sex effects on time-varying connectivity, with females exhibiting higher consistency than males. This observation may be attributed to the fact that many cognitive process, such as empathy for pain (Christov-Moore and Iacoboni, 2019; Groen et al., 2013) and language processing (Burman et al., 2008), typically involve higher levels of brain activation in females. However, due to the complex nature of the movie stimuli, it is difficult to pinpoint a specific cognitive process that can solely explain the observed sex differences. Moreover, sex differences may interact with other factors such as age (Etchell et al., 2018). Although we did not explore the interaction effects in the current study due to the limited sample size, it is important to investigate these effects in the future with larger samples.

In contrast to our initial hypothesis, the behavioral prediction analyses showed that only stationary connectivity, but not time-varying connectivity or regional activity, was able to predict FSIQ scores. Specifically, stationary connectivity demonstrated consistent prediction accuracy for FSIQ scores across both video clips. It is worth noting that the movie clips used in our current study primarily involved social interactions, which may not directly relate to general intelligence abilities. Previous research has shown that brain regions associated with FSIQ are typically higher-order association regions, such as the lateral prefrontal cortex (Cole et al., 2012; Geake and Hansen, 2005). The prediction features in the current analysis (Figure 5C and 5D) supported this point. Because stationary connectivity remains stable across different conditions, it may reflect general characteristics of cognitive functions, including FSIQ (Finn and Bandettini, 2021). On the other hand, time-varying connectivity, being more sensitive to certain movie contents, may not necessarily reflect general cognitive ability.

Regarding SCQ scores, none of the brain measures were able to predict individual differences in SCQ scores. The SCQ score was chosen because it reflects deficits in social functions related to autism. A previous study utilizing the same HBN dataset has reported an association between brain activity and SCQ scores at certain time points (Richardson, 2019). However, our whole-brain predictive modeling approach with cross-validation did not yield reliable associations between brain measures and SCQ scores. It is possible that such associations exist but may be limited to specific brain activation during certain events. Alternatively, in this normative sample, there may not have been a wide range of SCQ scores, which could weaken the association between brain measures and SCQ scores in a healthy sample.

The prediction accuracies of FSIQ using stationary connectivity, as measured by the correlation between actual and predicted cores, were on the order of 0.3. This level of accuracy is consistent with findings from previous studies on brain-behavior predictions (Cui and Gong, 2018; Di et al., 2022a; Marek et al., 2022). It is important to note that this effect size is reasonable considering the limitations in measurement reliability (Zuo et al., 2019). However, a correlation around 0.3 is usually considered moderate and suggests that the model can explain approximately 10% of the variance in the actual FSIQ measurements. This highlights the current bottleneck of the current brain-behavior prediction studies (Marek et al., 2022), which hiders the practical application of using brain imaging measures to accurately reflect behaviors. Further research is needed to enhance measurement accuracies and reliability in order to develop more robust models for brain-behavior predictions (Milham et al., 2021).

In this study, we analyzed two animated video clips: ‘The Present’ and a segment from the movie ‘Despicable Me.’ These clips differed in various aspects, including duration and content. ‘The Present’ is a self-constraint three-minute video, while the ‘Despicable Me’ segment is part of a longer one-hour movie. Due to the discrepancy in duration, the “Despicable Me” clip provided more time points for analysis, potentially resulting in more reliable connectivity measures.

Regarding brain connectivity, the averaged stationary connectivity matrices showed similarities between the two videos, suggesting consistent connectivity patterns. However, the time-varying connectivity patterns differed, which aligns with our previous findings (Di et al., 2022b). Interestingly, the preferred age and sex effects for both stationary connectivity and time-varying connectivity exhibited substantial differences between the two clips. This discrepancy may be partly attributed to thresholding effects, wherein reducing the threshold to reveal more effects made the age and sex effects appear more similar. However, some disparities still remained, indicating that individual differences may be more specific movie contents. While this variability in response to different movie stimuli can be advantageous, as they may target distinct brain functions, it also highlights the need for caution when interpreting results from movie watching studies. These studies are highly sensitive to the specific stimuli presented, emphasizing the necessity for careful analysis and interpretation.

The current study employed a data-driven ICA method to reduce the dimensionality of the fMRI data. Interestingly, our results revealed a unique component of the secondary sensorimotor regions, which differed from our previous work with other movie-watching data (Di et al., 2022b; Di and Biswal, 2022). This suggests that the data-driven approach is effective in extracting information specific to the sample and movie contents analyzed in the current study. Such specificity can be critical for some analysis. For example, when predicting FSIQ scores, the behavioral predictions were only significant when utilizing the data-driven ICA, as opposed to using the independent templates. However, it is worth noting that many results demonstrated consistent patterns when using the independent templates, compared to the data-driven ICA approach. For example, the majority of significant age effects exhibited age square or log age square effects, and females in general exhibited stronger stationary connectivity and time-varying connectivity than males. Their general patterns are robust across different data reduction schemes, providing further support for their reliability.

Another important consideration is the potential presence of sampling bias, which can impact the reliability and generalizability of the current findings. In this study, we implemented stringent exclusion criteria for MRI processing to ensure measurement quality, which may inadvertently introduce bias. However, we conducted analyses comparing the included and excluded samples and found no significant differences in terms of sex distributions, cognitive ability (FSIQ), and socioeconomic status (GSMSS). Nonetheless, it is worth noting that the excluded group was significantly younger than the included group, which could be a limitation that needs to be acknowledged. Furthermore, there may be other potential sources of sampling bias, such as the recruitment strategy and voluntary participation of participants (Alexander et al., 2017). These factors can influence the composition of the sample and potentially limit the generalizability of the findings. Therefore, it is important to confirm the generalizability of the results through further studies that include diverse populations and employ rigorous sampling methods.

## 5. Conclusion

In the current study, we examined age and sex effects, and behavioral correlates of time-varying and stationary connectivity during movie watching. We found that time-varying connectivity exhibited greater sensitivity to the age and sex effects compared with stationary connectivity. However, regarding predicting individuals’ FSIQ scores, only stationary connectivity demonstrated predictive capabilities.

These results provide a more comprehensive understanding of the individual differences observed in time-varying and stationary connectivity within the human brain.

## Acknowledgement

This study was supported by (US) National Institute of Mental Health grants to X.D. (R15MH125332) and B.B.B. (R01MH131335). The authors would like to thank Donna Chen for her comments on earlier versions of this manuscript.

## Reference

Abrol, A., Damaraju, E., Miller, R.L., Stephen, J.M., Claus, E.D., Mayer, A.R., Calhoun, V.D., 2017. Replicability of time-varying connectivity patterns in large resting state fMRI samples. NeuroImage 163, 160–176. https://doi.org/10.1016/j.neuroimage.2017.09.020

Alexander, L.M., Escalera, J., Ai, L., Andreotti, C., Febre, K., Mangone, A., Vega-Potler, N., Langer, N., Alexander, A., Kovacs, M., Litke, S., O’Hagan, B., Andersen, J., Bronstein, B., Bui, A., Bushey, M., Butler, H., Castagna, V., Camacho, N., Chan, E., Citera, D., Clucas, J., Cohen, S., Dufek, S., Eaves, M., Fradera, B., Gardner, J., Grant-Villegas, N., Green, G., Gregory, C., Hart, E., Harris, S., Horton, M., Kahn, D., Kabotyanski, K., Karmel, B., Kelly, S.P., Kleinman, K., Koo, B., Kramer, E., Lennon, E., Lord, C., Mantello, G., Margolis, A., Merikangas, K.R., Milham, J., Minniti, G., Neuhaus, R., Levine, A., Osman, Y., Parra, L.C., Pugh, K.R., Racanello, A., Restrepo, A., Saltzman, T., Septimus, B., Tobe, R., Waltz, R., Williams, A., Yeo, A., Castellanos, F.X., Klein, A., Paus, T., Leventhal, B.L., Craddock, R.C., Koplewicz, H.S., Milham, M.P., 2017. An open resource for transdiagnostic research in pediatric mental health and learning disorders. Scientific Data 4, 1–26. https://doi.org/10.1038/sdata.2017.181

Allen, E.A., Damaraju, E., Plis, S.M., Erhardt, E.B., Eichele, T., Calhoun, V.D., 2014. Tracking whole-brain connectivity dynamics in the resting state. Cerebral cortex (New York, N.Y. : 1991) 24, 663–76. https://doi.org/10.1093/cercor/bhs352

Allen, M., Frank, D., Glen, J.C., Fardo, F., Callaghan, M.F., Rees, G., 2017. Insula and somatosensory cortical myelination and iron markers underlie individual differences in empathy. Sci Rep 7, 43316. https://doi.org/10.1038/srep43316

Ashburner, J., 2007. A fast diffeomorphic image registration algorithm. NeuroImage 38, 95–113. https://doi.org/10.1016/j.neuroimage.2007.07.007

Biswal, B., Yetkin, F.Z., Haughton, V.M., Hyde, J.S., 1995. Functional connectivity in the motor cortex of resting human brain using echo-planar MRI. Magnetic resonance in medicine : official journal of the Society of Magnetic Resonance in Medicine / Society of Magnetic Resonance in Medicine 34, 537–41. https://doi.org/10.1002/mrm.1910340409

Biswal, B.B., Mennes, M., Zuo, X.-N., Gohel, S., Kelly, C., Smith, S.M., Beckmann, C.F., Adelstein, J.S., Buckner, R.L., Colcombe, S., Dogonowski, A.-M., Ernst, M., Fair, D., Hampson, M., Hoptman, M.J., Hyde, J.S., Kiviniemi, V.J., Kötter, R., Li, S.-J., Lin, C.-P., Lowe, M.J., Mackay, C., Madden, D.J., Madsen, K.H., Margulies, D.S., Mayberg, H.S., McMahon, K., Monk, C.S., Mostofsky, S.H., Nagel, B.J., Pekar, J.J., Peltier, S.J., Petersen, S.E., Riedl, V., Rombouts, S.A.R.B., Rypma, B., Schlaggar, B.L., Schmidt, S., Seidler, R.D., Siegle, G.J., Sorg, C., Teng, G.-J., Veijola, J., Villringer, A., Walter, M., Wang, L., Weng, X.-C., Whitfield-Gabrieli, S., Williamson, P., Windischberger, C., Zang, Y.-F., Zhang, H.-Y., Castellanos, F.X., Milham, M.P., 2010. Toward discovery science of human brain function. Proceedings of the National Academy of Sciences of the United States of America 107, 4734–9. https://doi.org/10.1073/pnas.0911855107

Burman, D.D., Bitan, T., Booth, J.R., 2008. Sex differences in neural processing of language among children. Neuropsychologia 46, 1349–1362. https://doi.org/10.1016/j.neuropsychologia.2007.12.021

Calhoun, V.D., Adali, T., Pearlson, G.D., Pekar, J.J., 2001. A method for making group inferences from functional MRI data using independent component analysis. Human brain mapping 14, 140–51.

Cantlon, J.F., Li, R., 2013. Neural Activity during Natural Viewing of Sesame Street Statistically Predicts Test Scores in Early Childhood. PLOS Biology 11, e1001462. https://doi.org/10.1371/journal.pbio.1001462

Cao, M., Wang, J.H., Dai, Z.J., Cao, X.Y., Jiang, L.L., Fan, F.M., Song, X.W., Xia, M.R., Shu, N., Dong, Q., Milham, M.P., Castellanos, F.X., Zuo, X.N., He, Y., 2014. Topological organization of the human brain functional connectome across the lifespan. Developmental Cognitive Neuroscience 7, 76–93. https://doi.org/10.1016/j.dcn.2013.11.004

Christov-Moore, L., Iacoboni, M., 2019. Sex differences in somatomotor representations of others’ pain: a permutation-based analysis. Brain Struct Funct 224, 937–947. https://doi.org/10.1007/s00429-018-1814-y

Cohen, S.S., Tottenham, N., Baldassano, C., 2022. Developmental changes in story-evoked responses in the neocortex and hippocampus. eLife 11, e69430. https://doi.org/10.7554/eLife.69430

Cole, M.W., Yarkoni, T., Repovš, G., Anticevic, A., Braver, T.S., 2012. Global Connectivity of Prefrontal Cortex Predicts Cognitive Control and Intelligence. J. Neurosci. 32, 8988–8999. https://doi.org/10.1523/JNEUROSCI.0536-12.2012

Cui, Z., Gong, G., 2018. The effect of machine learning regression algorithms and sample size on individualized behavioral prediction with functional connectivity features. NeuroImage. https://doi.org/10.1016/j.neuroimage.2018.06.001

Di, X., Biswal, B.B., 2023. A functional MRI pre-processing and quality control protocol based on statistical parametric mapping (SPM) and MATLAB. Frontiers in Neuroimaging 1.

Di, X., Biswal, B.B., 2022. Principal component analysis reveals multiple consistent responses to naturalistic stimuli in children and adults. Human Brain Mapping 43, 3332–3345. https://doi.org/10.1002/hbm.25568

Di, X., Biswal, B.B., 2020. Intersubject consistent dynamic connectivity during natural vision revealed by functional MRI. NeuroImage 116698. https://doi.org/10.1016/j.neuroimage.2020.116698

Di, X., Woelfer, M., Kühn, S., Zhang, Z., Biswal, B.B., 2022a. Estimations of the weather effects on brain functions using functional MRI: A cautionary note. Human Brain Mapping 43, 3346–3356. https://doi.org/10.1002/hbm.25576

Di, X., Zhang, Z., Xu, T., Biswal, B.B., 2022b. Dynamic and stationary brain connectivity during movie watching as revealed by functional MRI. Brain Struct Funct. https://doi.org/10.1007/s00429-022-02522-w

Eichenbaum, A., Pappas, I., Lurie, D., Cohen, J.R., D’Esposito, M., 2021. Differential contributions of static and time-varying functional connectivity to human behavior. Network Neuroscience 5, 145–165. https://doi.org/10.1162/netn_a_00172

Etchell, A., Adhikari, A., Weinberg, L.S., Choo, A.L., Garnett, E.O., Chow, H.M., Chang, S.-E., 2018. A systematic literature review of sex differences in childhood language and brain development. Neuropsychologia 114, 19–31. https://doi.org/10.1016/j.neuropsychologia.2018.04.011

Faghiri, A., Stephen, J.M., Wang, Y.-P., Wilson, T.W., Calhoun, V.D., 2018. Changing brain connectivity dynamics: From early childhood to adulthood. Human Brain Mapping 39, 1108–1117. https://doi.org/10.1002/hbm.23896

Faskowitz, J., Esfahlani, F.Z., Jo, Y., Sporns, O., Betzel, R.F., 2020. Edge-centric functional network representations of human cerebral cortex reveal overlapping system-level architecture. Nat Neurosci 23, 1644–1654. https://doi.org/10.1038/s41593-020-00719-y

Finn, E.S., Bandettini, P.A., 2021. Movie-watching outperforms rest for functional connectivity-based prediction of behavior. NeuroImage 235, 117963. https://doi.org/10.1016/j.neuroimage.2021.117963

Friston, K.J., 1994. Functional and effective connectivity in neuroimaging: A synthesis. Human Brain Mapping 2, 56–78. https://doi.org/10.1002/hbm.460020107

Friston, K.J., Williams, S., Howard, R., Frackowiak, R.S., Turner, R., 1996. Movement-related effects in fMRI time-series. Magnetic resonance in medicine : official journal of the Society of Magnetic Resonance in Medicine / Society of Magnetic Resonance in Medicine 35, 346–55. https://doi.org/DOI 10.1002/mrm.1910350312

Fu, Z., Tu, Y., Di, X., Du, Y., Sui, J., Biswal, B.B., Zhang, Z., de Lacy, N., Calhoun, V.D., 2019. Transient increased thalamic-sensory connectivity and decreased whole-brain dynamism in autism. NeuroImage 190, 191–204. https://doi.org/10.1016/j.neuroimage.2018.06.003

Geake, J.G., Hansen, P.C., 2005. Neural correlates of intelligence as revealed by fMRI of fluid analogies. NeuroImage 26, 555–564. https://doi.org/10.1016/j.neuroimage.2005.01.035

Groen, Y., Wijers, A.A., Tucha, O., Althaus, M., 2013. Are there sex differences in ERPs related to processing empathy-evoking pictures? Neuropsychologia 51, 142–155. https://doi.org/10.1016/j.neuropsychologia.2012.11.012

Hasson, U., Nir, Y., Levy, I., Fuhrmann, G., Malach, R., 2004. Intersubject synchronization of cortical activity during natural vision. Science (New York, N.Y.) 303, 1634–40. https://doi.org/10.1126/science.1089506

Jin, C., Jia, H., Lanka, P., Rangaprakash, D., Li, L., Liu, T., Hu, X., Deshpande, G., 2017. Dynamic brain connectivity is a better predictor of PTSD than static connectivity. Human Brain Mapping 38, 4479–4496. https://doi.org/10.1002/hbm.23676

Kauppi, J.-P., Jääskeläinen, I.P., Sams, M., Tohka, J., 2010. Inter-subject correlation of brain hemodynamic responses during watching a movie: localization in space and frequency. Front. Neuroinform. 4. https://doi.org/10.3389/fninf.2010.00005

Lamm, C., Nusbaum, H.C., Meltzoff, A.N., Decety, J., 2007. What Are You Feeling? Using Functional Magnetic Resonance Imaging to Assess the Modulation of Sensory and Affective Responses during Empathy for Pain. PLOS ONE 2, e1292. https://doi.org/10.1371/journal.pone.0001292

Marek, S., Tervo-Clemmens, B., Calabro, F.J., Montez, D.F., Kay, B.P., Hatoum, A.S., Donohue, M.R., Foran, W., Miller, R.L., Hendrickson, T.J., Malone, S.M., Kandala, S., Feczko, E., Miranda-Dominguez, O., Graham, A.M., Earl, E.A., Perrone, A.J., Cordova, M., Doyle, O., Moore, L.A., Conan, G.M., Uriarte, J., Snider, K., Lynch, B.J., Wilgenbusch, J.C., Pengo, T., Tam, A., Chen, J., Newbold, D.J., Zheng, A., Seider, N.A., Van, A.N., Metoki, A., Chauvin, R.J., Laumann, T.O., Greene, D.J., Petersen, S.E., Garavan, H., Thompson, W.K., Nichols, T.E., Yeo, B.T.T., Barch, D.M., Luna, B., Fair, D.A., Dosenbach, N.U.F., 2022. Reproducible brain-wide association studies require thousands of individuals. Nature 603, 654–660. https://doi.org/10.1038/s41586-022-04492-9

Margulies, D.S., Ghosh, S.S., Goulas, A., Falkiewicz, M., Huntenburg, J.M., Langs, G., Bezgin, G., Eickhoff, S.B., Castellanos, F.X., Petrides, M., Jefferies, E., Smallwood, J., 2016. Situating the default-mode network along a principal gradient of macroscale cortical organization. PNAS 113, 12574–12579. https://doi.org/10.1073/pnas.1608282113

Marusak, H.A., Calhoun, V.D., Brown, S., Crespo, L.M., Sala-Hamrick, K., Gotlib, I.H., Thomason, M.E., 2017. Dynamic functional connectivity of neurocognitive networks in children. Human Brain Mapping 38, 97–108. https://doi.org/10.1002/hbm.23346

Milham, M.P., Vogelstein, J., Xu, T., 2021. Removing the Reliability Bottleneck in Functional Magnetic Resonance Imaging Research to Achieve Clinical Utility. JAMA Psychiatry 78, 587–588. https://doi.org/10.1001/jamapsychiatry.2020.4272

Nastase, S.A., Gazzola, V., Hasson, U., Keysers, C., 2019. Measuring shared responses across subjects using intersubject correlation. Soc Cogn Affect Neurosci 14, 667–685. https://doi.org/10.1093/scan/nsz037

Nickerson, L.D., Smith, S.M., Öngür, D., Beckmann, C.F., 2017. Using Dual Regression to Investigate Network Shape and Amplitude in Functional Connectivity Analyses. Frontiers in Neuroscience 11.

O’Connor, D., Potler, N.V., Kovacs, M., Xu, T., Ai, L., Pellman, J., Vanderwal, T., Parra, L.C., Cohen, S., Ghosh, S., Escalera, J., Grant-Villegas, N., Osman, Y., Bui, A., Craddock, R.C., Milham, M.P., 2017. The Healthy Brain Network Serial Scanning Initiative: a resource for evaluating inter-individual differences and their reliabilities across scan conditions and sessions. Gigascience 6, 1–14. https://doi.org/10.1093/gigascience/giw011

Petroni, A., Cohen, S.S., Ai, L., Langer, N., Henin, S., Vanderwal, T., Milham, M.P., Parra, L.C., 2018. The Variability of Neural Responses to Naturalistic Videos Change with Age and Sex. eNeuro 5. https://doi.org/10.1523/ENEURO.0244-17.2017

Portet, S., 2020. A primer on model selection using the Akaike Information Criterion. Infectious Disease Modelling 5, 111–128. https://doi.org/10.1016/j.idm.2019.12.010

Rashid, B., Arbabshirani, M.R., Damaraju, E., Cetin, M.S., Miller, R., Pearlson, G.D., Calhoun, V.D., 2016. Classification of schizophrenia and bipolar patients using static and dynamic resting-state fMRI brain connectivity. NeuroImage 134, 645–657. https://doi.org/10.1016/j.neuroimage.2016.04.051

Rashid, B., Blanken, L.M.E., Muetzel, R.L., Miller, R., Damaraju, E., Arbabshirani, M.R., Erhardt, E.B., Verhulst, F.C., van der Lugt, A., Jaddoe, V.W.V., Tiemeier, H., White, T., Calhoun, V., 2018. Connectivity dynamics in typical development and its relationship to autistic traits and autism spectrum disorder. Human Brain Mapping 39, 3127–3142. https://doi.org/10.1002/hbm.24064

Richardson, H., 2019. Development of brain networks for social functions: Confirmatory analyses in a large open source dataset. Developmental Cognitive Neuroscience 37, 100598. https://doi.org/10.1016/j.dcn.2018.11.002

Rutter, M., Bailey, A., Lord, C., 2003. The social communication questionnaire.

Tian, L., Ye, M., Chen, C., Cao, X., Shen, T., 2021. Consistency of functional connectivity across different movies. NeuroImage 233, 117926. https://doi.org/10.1016/j.neuroimage.2021.117926

Vanderwal, T., Eilbott, J., Castellanos, F.X., 2019. Movies in the magnet: Naturalistic paradigms in developmental functional neuroimaging. Developmental Cognitive Neuroscience 36, 100600. https://doi.org/10.1016/j.dcn.2018.10.004

Vanderwal, T., Eilbott, J., Kelly, C., Frew, S.R., Woodward, T.S., Milham, M.P., Castellanos, F.X., 2021. Stability and similarity of the pediatric connectome as developmental measures. NeuroImage 226, 117537. https://doi.org/10.1016/j.neuroimage.2020.117537

Vieira, B.H., Pamplona, G.S.P., Fachinello, K., Silva, A.K., Foss, M.P., Salmon, C.E.G., 2022. On the prediction of human intelligence from neuroimaging: A systematic review of methods and reporting. Intelligence 93, 101654. https://doi.org/10.1016/j.intell.2022.101654

Wagenmakers, E.-J., Farrell, S., 2004. AIC model selection using Akaike weights. Psychonomic Bulletin & Review 11, 192–196. https://doi.org/10.3758/BF03206482

Wechsler, D., 2014. The Wechsler intelligence scale for children—fifth edition.

Xia, M., Wang, J., He, Y., 2013. BrainNet Viewer: a network visualization tool for human brain connectomics. PloS one 8, e68910. https://doi.org/10.1371/journal.pone.0068910

Yarkoni, T., Poldrack, R.A., Nichols, T.E., Van Essen, D.C., Wager, T.D., 2011. Large-scale automated synthesis of human functional neuroimaging data. Nature methods 8, 665–70. https://doi.org/10.1038/nmeth.1635

Yeo, B.T.T., Krienen, F.M., Sepulcre, J., Sabuncu, M.R., Lashkari, D., Hollinshead, M., Roffman, J.L., Smoller, J.W., Zöllei, L., Polimeni, J.R., Fischl, B., Liu, H., Buckner, R.L., 2011. The organization of the human cerebral cortex estimated by intrinsic functional connectivity. Journal of neurophysiology 106, 1125–65. https://doi.org/10.1152/jn.00338.2011

Zuo, X.-N., Kelly, C., Di Martino, A., Mennes, M., Margulies, D.S., Bangaru, S., Grzadzinski, R., Evans, A.C., Zang, Y.-F., Castellanos, F.X., Milham, M.P., 2010. Growing together and growing apart: regional and sex differences in the lifespan developmental trajectories of functional homotopy. The Journal of neuroscience : the official journal of the Society for Neuroscience 30, 15034–43. https://doi.org/10.1523/JNEUROSCI.2612-10.2010

Zuo, X.-N., Xu, T., Milham, M.P., 2019. Harnessing reliability for neuroscience research. Nature Human Behaviour 1. https://doi.org/10.1038/s41562-019-0655-x

