## Supplementary Materials for "Individual differences in time-varying and stationary brain connectivity during movie watching from childhood to early adulthood: age, sex, and behavioral associations"

This file contains Supplementary Figures S1 to S8.

**Figure S1.** Sample exclusion and inclusion flowchart

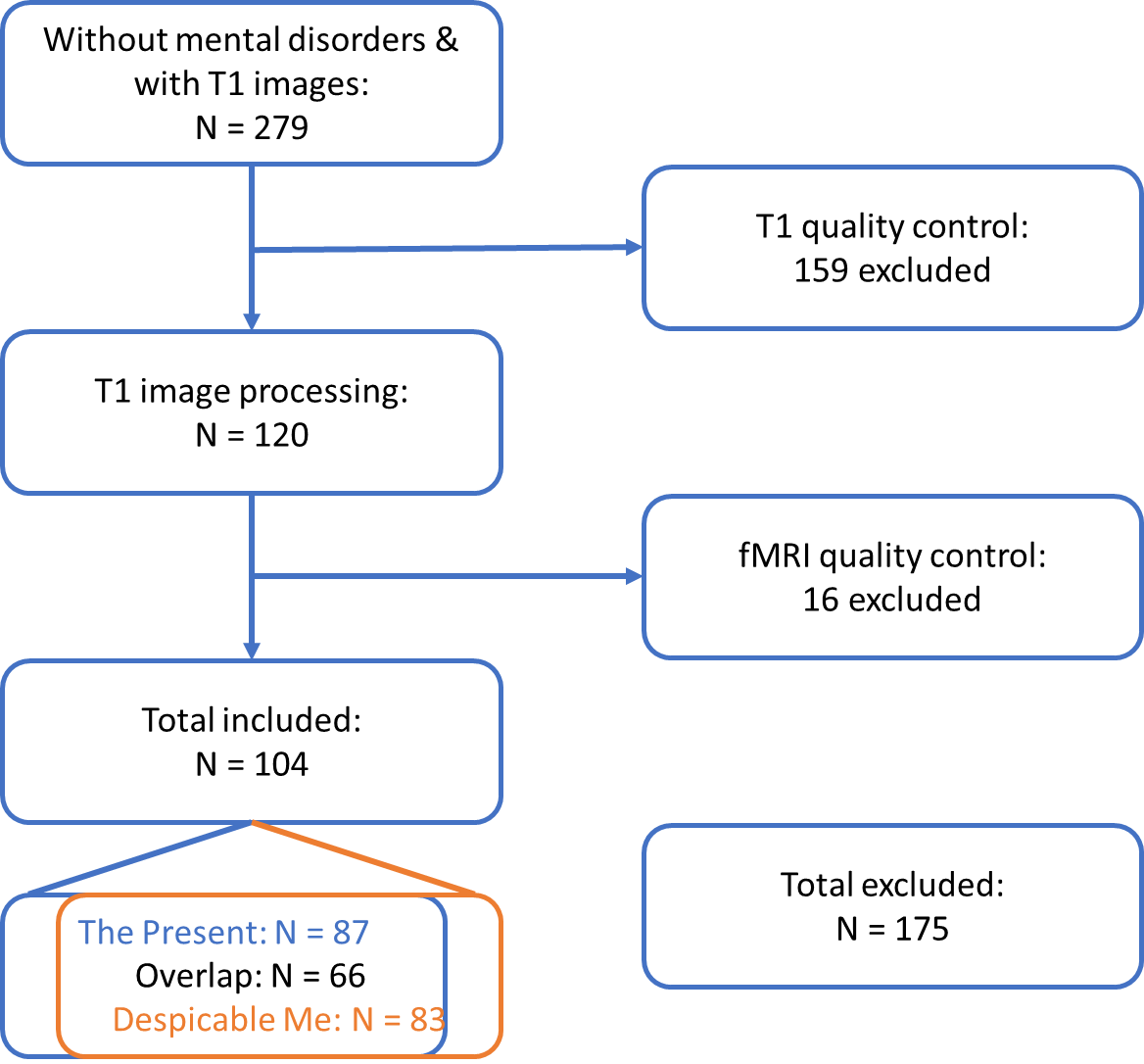

**Figure S2. Effects of head motion**

A, Inter-individual consistency of head motion (percent variance explained by the first principal component of framewise displacement (FD) time series). B, model comparison results of different age models for mean FD. For the clip of The Present, the constant models were slightly favorable for both translation and rotation, indicating no age effects. For the clip of Despicable Me, the log age models were slightly favorable for both directions. C through F show the scatter plots of mean FD for both directions and clips with fitted age curves.

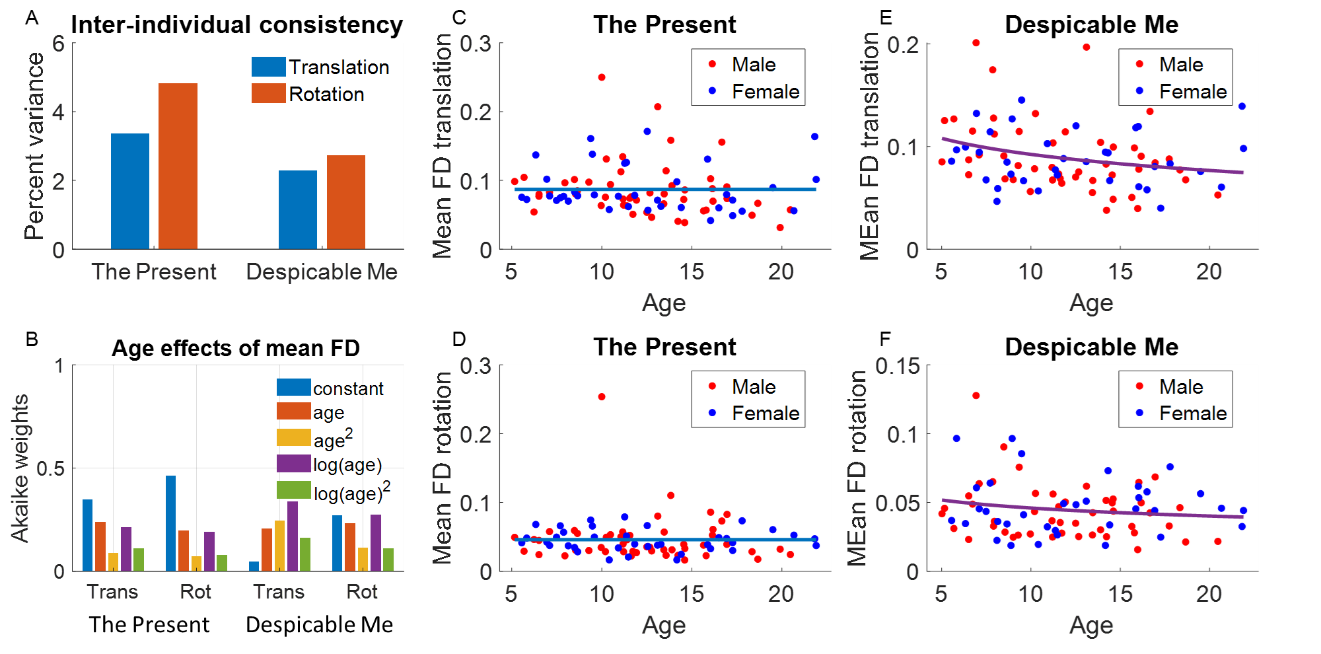

**Figure S3.** Illustration of the behavioral prediction analysis.

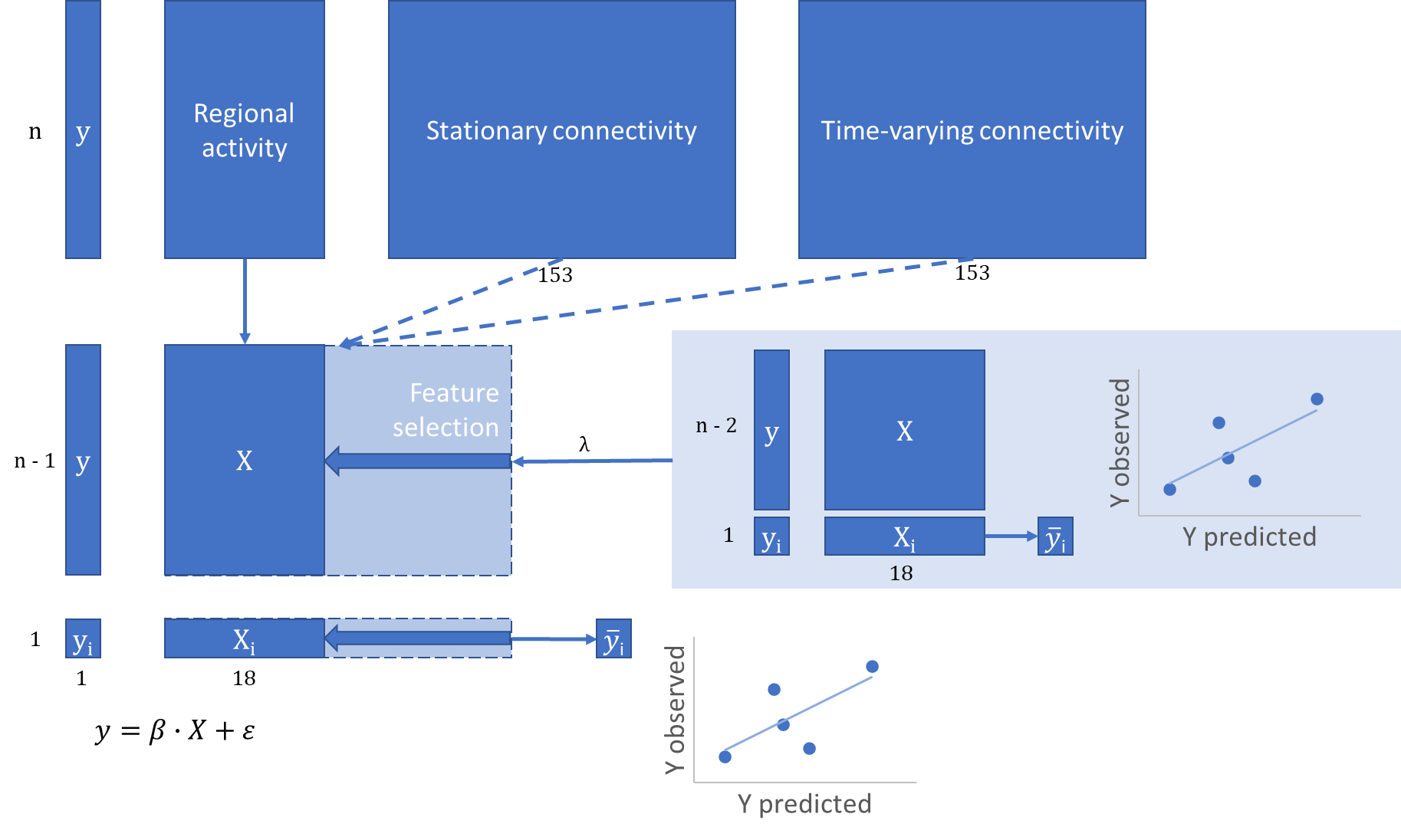

**Figure S4.** Model comparison results for the stationary connectivity

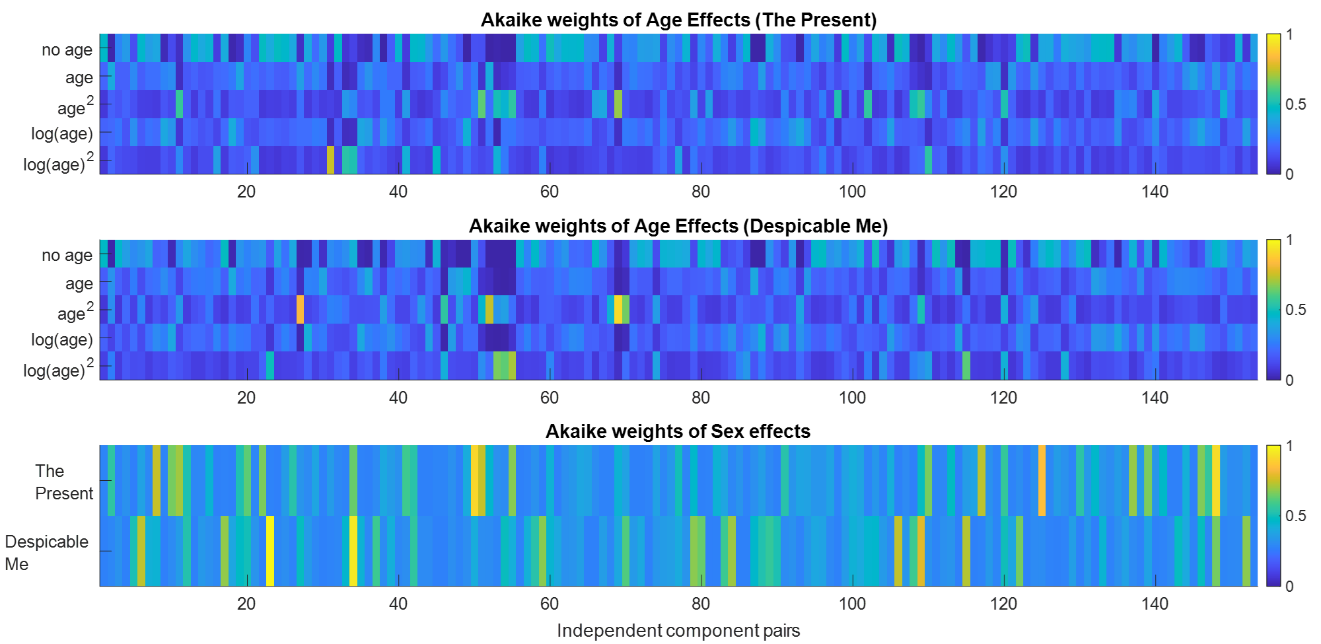

**Figure S5.** Model comparison results for the time-varying connectivity

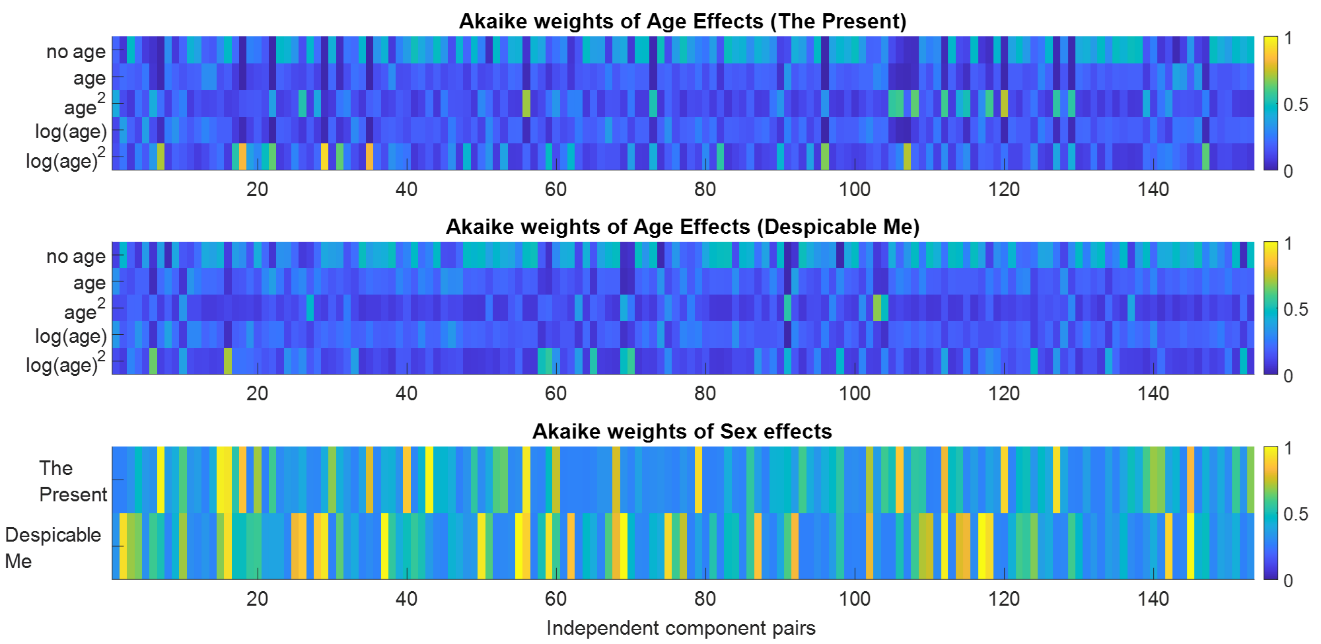

**Figure S6.** Machine learning prediction results for predicting full-scale intelligence quotients (FSIQ).

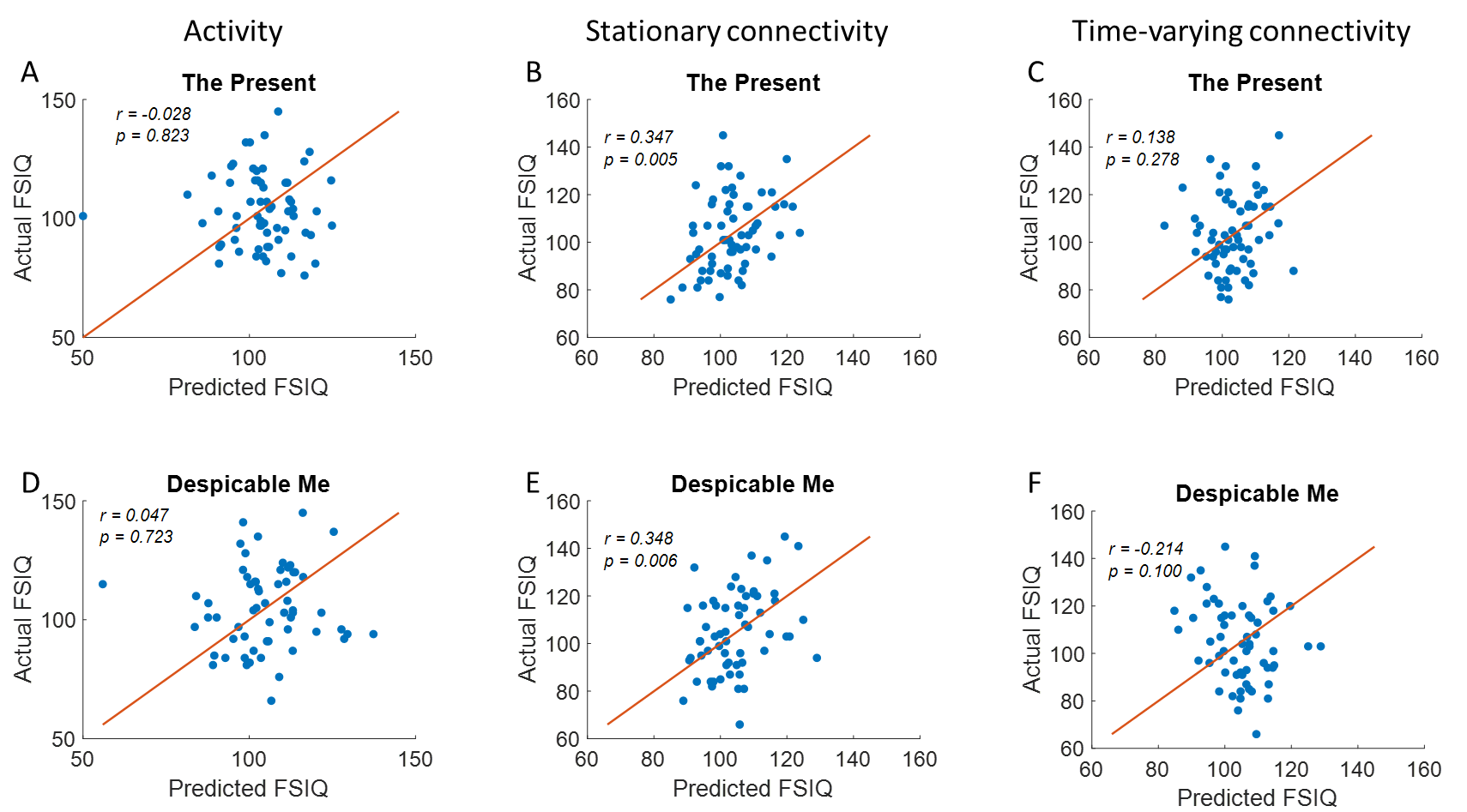

**Figure S7.** Machine learning prediction results for predicting the SCQ scores (Social Communication Questionnaire).

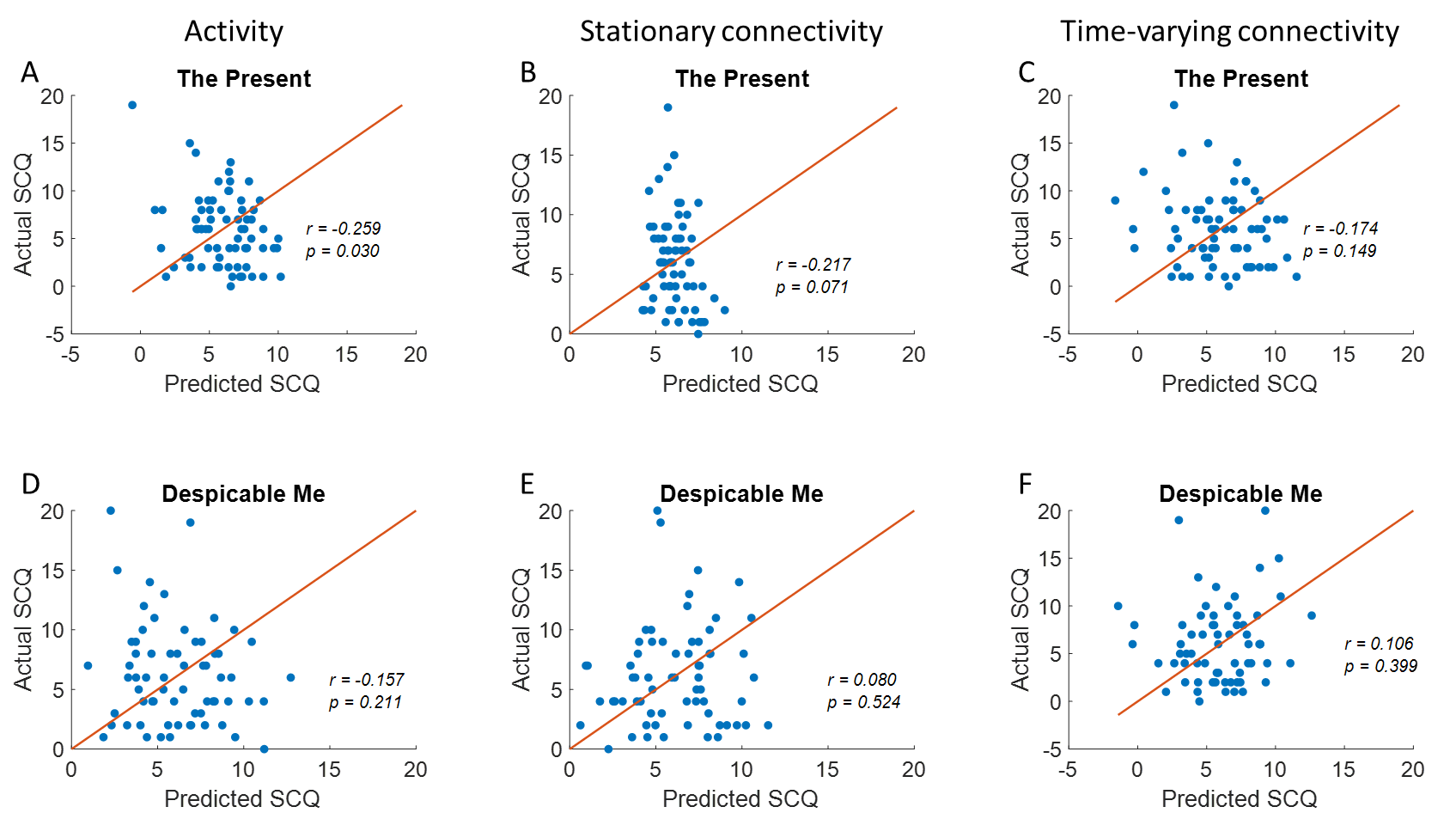

**Figure S8** A, Representative maps of 16 independent components from Di et al., (2022). B, Inter-individual consistency of regional activity (percent variance explained by the first principal component) for the two video clips. C, D, and E, Model comparison results for different age models and sex effects for regional activity for the two video clips.

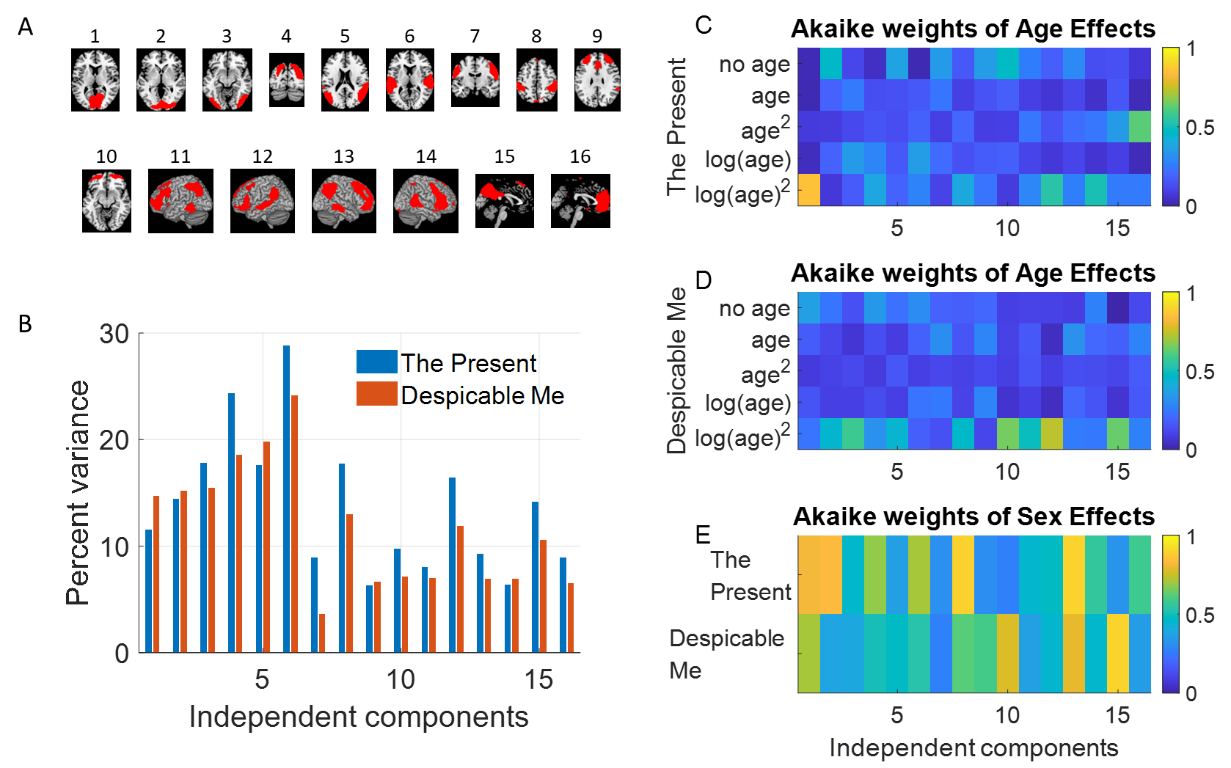
